# Content of xylose, trehalose and L-citrulline in cucumber fermentations and utilization of such compounds by certain lactobacilli

**DOI:** 10.1101/2019.12.20.885608

**Authors:** Redife Aslihan Ucar, Ilenys M. Pérez-Díaz, Lisa L. Dean

## Abstract

This research determined the concentration of trehalose, xylose and L-citrulline in fresh and fermented cucumbers and their utilization by *Lactobacillus pentosus*, *Lactobacillus plantarum*, *Lactobacillus brevis* and *Lactobacillus buchneri*. Targeted compounds were measured by HPLC and the ability of certain lactobacilli to utilize them was scrutinized in fermented cucumber juice. Fresh cucumber juice was supplemented with trehalose, xylose and L-citrulline to observed mixed culture fermentations. Changes in the biochemistry, pH and colony counts during fermentations were monitored. Trehalose was detected in fermentations to 15.51 ± 1.68 mM. Xylose was found in a fresh cucumber sample to 36.05 mM. L-citrulline was present in fresh and fermented cucumber samples to 1.05 ± 0.63 mM. Most of the lactobacilli tested utilized trehalose and xylose in FCJM at pH 4.7. L-citrulline was utilized by *L. buchneri* and produced by other LAB. L-citrulline (12.43 ± 2.3 mM) was converted to ammonia (14.54 ± 3.60 mM) and the biogenic amine ornithine (14.19 ± 1.07 mM) by *L. buchneri* at pH 4.7 in the presence of 0.5 ± 0.2 mM glucose enhancing growth by 0.5 log CFU/mL. The use of a mixed starter culture containing *L. buchneri* aided in the removal of L-citrulline and enhanced the fermentation stability. The utilization of L-citrulline by *L. buchneri* may be a cause of concern for the stability of cucumber fermentations at pH 3.7 or above. This study identifies the use of a tripartite starter culture as an enhancer of microbial stability for fermented cucumbers.

**Importance:** Utilization of trehalose, xylose and L-citrulline as energy sources by the indigenous cucumber microbiota was studied to determine if this was a cause for the spoilage of the fermented fruit. While the plant derived sugars, trehalose and xylose, were occasionally present in cucumber fermentations, they are readily utilized by the bacteria spearheading primary fermentation. L-citrulline, however, is an amino acid not naturally found in proteins, was detected in all the samples tested and was uniquely utilized by the spoilage associated bacterium, *L. buchneri*. Additionally, the bacteria involved in cucumber fermentation produced L-citrulline. It was observed that the use of *L. pentosus*, *L. brevis* and *L. buchneri* in a mixed starter culture aids in the removal of the alternate energy sources, including L-citrulline, and the generation of a stable cucumber fermentation for 55 days under anaerobiosis.

## 1. Introduction

The utilization of glucose and fructose by *Lactobacillus pentosus* and *Lactobacillus plantarum* is the main biochemical conversion in cucumber fermentation. The transformation of sugars to lactic and acetic acids is typically monitored using HPLC and changes in pH in research laboratories and the industrial setting, respectively. Other compounds that could serve as energy sources for microbes are present in fresh and fermented cucumbers. Sucrose and xylose were extracted from cucumber mesocarp and endocarp (1, 2, 3). Rhamnose, fucose, arabinose, mannose, galactose, and galacturonic acid have been extracted from cucumber mesocarp at concentrations between 0.1 and 7 mM and found not to change as a function of tissue softening induced by heating or fermentation (2). The concentrations of L-citrulline, trehalose, cellobiose, xylose, lyxose, gentiobiose, furfural, and lactic acid were found to change in anaerobic fermented cucumber spoilage by *L. buchneri* using two-dimensional time of flight mass spectrometry (4). L-citrulline, D-trehalose, and D-cellobiose are utilized by *L. buchneri* prior to fermented cucumber spoilage characterized by lactic acid degradation (4). Cellobiose, trehalose, and gentiobiose have also been found in traditional pickles in Sichuan, China (5) and in the traditional Indian vegetable fermented products gundruk, sinki, and kalphi (6). It has been postulated that the utilization of alternative energy sources in cucumber fermentation supports the growth and metabolic activity of spoilage associated microbes such as *L. buchneri* (4).

The lactobacilli typically prevailing in cucumber fermentations such as *L. pentosus* and *L. plantarum* are notorious for their ability to utilize a variety of carbon sources (7, 8, 9). Lactobacilli species have been used as functional starter cultures for fermented food products such as plant-derived foods, meats, wine, cheese, beer and sourdough. *L. plantarum* NCIMB 8026 is able to utilize ribose, xylose and L-arabinose in the presence of glucose (10). A group of 185 *L. plantarum* strains was found to utilize D-trehalose (11). A substantial number of *Lactobacillus casei, L. plantarum*, *Lactobacillus buchneri* and *Lactobacillus brevis* strains are able to grow on D-trehalose and D-xylose (12).

We hypothesize that utilization of alternative energy sources by the organisms prevailing in cucumber fermentations, including *L. pentosus*, *L. plantarum* and *L. brevis* hampers the ability of spoilage-associated microbes, such as *L. buchneri*, to derive energy for growth and/or metabolic activity post-fermentation. In a previous study, we observed that while the disaccharides cellobiose and gentiobiose can be utilized by several lactic acid bacteria (LAB) in fermented cucumber juice (FCJM) at pH 4.7, the concentration of such disaccharides that is freely available in fresh and fermented cucumbers is less than 10 µM (13). This study evaluates the intrinsic concentration of trehalose, xylose and L-citrulline in fresh and fermented cucumbers and determines the ability of certain LAB to utilize such compounds in FCJM to simulate conditions post-primary fermentation.

Trehalose is a disaccharide composed of two glucose units joined by an α-α, (1,1) linkage produced by organisms (14). Although, some non-lactic acid producing microbes such as *Escherichia coli* and *Saccharomyces cerevisiae* can synthesize trehalose, most of the literature recognizes the ability of a wider spectrum of microbes capable of transporting and utilizing the disaccharide (15). Trehalose is known to accumulate up to 12% of the plant dry weight, in cryptobiotic species, and impart stress tolerance, particularly drought (16). Bacteria that are freeze-dried in the presence of trehalose recover significantly better that those treated with sucrose and retained viability even after extended exposure to high humidity (14, 17). Given that trehalose lacks reducing ends, it is resistant to heat, extreme pH and the Maillard’s reaction, and stabilizes biological structures under stress (16). Trehalose forms a glass-like structure around biological membranes and enzymes after dehydration (18).

Xylose, a pentose sugar, is metabolized by several LAB via the Phosphoketolase Pathway and, in some cases, via the Embden-Meyerhof Pathway (19, 20, 21, 22, 23, 5, 24). The pentose is transported by *L. pentosus* via a phosphotransferase system involving the enzymatic products of *xylP* and *xylQ* through the EII^MAN^ transporter system (23). Proteomic analysis of *L. brevis* suggests the metabolism of xylose alone or concomitantly with glucose to proceed heterofermentatively (25, 5). *L. brevis* cells supplemented with xylose alone or glucose and xylose express the *xyl* operon (25). However, *L. plantarum* strains able to ferment pentoses have been found unable to grow in culture media supplemented with xylose (26, 10). A group of 185 *L. plantarum* strains were found to utilize D-trehalose, but not xylose (11). Xylose utilization by *L. plantarum* has been observed by co-inoculating two or more strains in MRS supplemented with the sugar (11). It is speculated that carbohydrates may be co-metabolized by *L. plantarum* with different purposes.

L-citrulline is a non-protein α-amino acid first isolated from watermelon in 1914 by Koga and Ohtake and identified in 1930 by Wada (27). The physiological level of L-citrulline in cantaloupe, cucumbers and pumpkin is 22.23 ± 3.69 mM (28). L-citrulline is involved in the detoxification of catabolic ammonia, in the production of the vasodilator, nitric oxide, and the precursor of arginine in the kidney of mammalians (28). In plants, L-citrulline has been associated with protection against oxidative stress, particularly during periods of drought (29). L-citrulline can be used as an alternate source for the generation of ATP by bacteria when fermentable carbohydrates are insufficient in the environment (30, 31). Some LAB produce L-citrulline from the degradation of arginine allowing the α-amino acid produced to be consumed by other lactobacilli (32). In the conversion of L-arginine and water to L-citrulline and ammonia via arginine deiminase (*arcA*), L-citrulline is phosphorylated to form L-ornithine and carbamyl phosphate via the ornithine transcarbamylase (*arcB*). Carbamyl phosphate and ADP react to form ATP, carbon dioxide and ammonia via the carbamate kinase (*arcC*). Thus, the catabolism of arginine can be used by LAB to derive energy in the form of ATP, particularly in the absence of sugars in acidic environments (33, 34, 30, 31). The conversion of ornithine into the biogenic amine putrescine and L-citrulline and/or carbamayl-phosphate to ethyl-carbamate, a carcinogen, in the presence of ethanol are two of the undesired consequences of arginine catabolism in fermented foods. However, the production of ammonia induces an increase in the extracellular pH (35, 33, 30).

In this study, we determined the concentrations of trehalose, xylose and L-citrulline in fresh cucumbers and cucumber fermentations using HPLC analysis. The ability of *L. plantarum*, *L. pentosus*, *L. brevis*, and *L. buchneri* to utilize trehalose, xylose and L-citrulline was also determined using FCJM to mimic conditions at the end of cucumber fermentation. Furthermore, the ability of *L. buchneri* to utilize L-citrulline in the presence and absence of glucose was conducted. To better understand the interactions of the LAB in cucumber fermentations we observed the changes in the biochemistry, pH and colony counts in co-cultures of LAB inoculated in fresh cucumber juice (FrCJ) medium and FCJM supplemented with xylose, trehalose and L-citrulline.

## 2. Materials and methods

### 2.1. Preparation of fresh and fermented cucumber samples for HPLC analysis

Samples of four fresh, size 2B, pickling cucumber lots to be fermented in commercial vessels were obtained from a local processor. The corresponding fermented cucumber samples were collected on days 3 and 38 of fermentation together with the fermentation cover brine in a 50:50 ratio. Fresh and fermented cucumbers were sliced using aseptic techniques and blended for 90 s at medium speed using a Waring Commercial Blender 700S (Torrington, CT, USA) equipped with sterilized glass cups. Fermented cucumbers were blended together with the fermentation cover brine in a 50:50 ratio. Cucumber slurries were homogenized using a Seward Stomacher 400 (Bohemia, NY, USA) in 6” x 4.5” filter stomacher bags for 1 min at maximum speed. One mL aliquots of the filtered homogenate were spun at 15,294 rcf for 10 min in an Eppendorf benchtop refrigerated centrifuge 5810R (Hamburg, Germany) to remove residual particulate matter. Supernatants were used for HPLC analyses conducted as described below.

### 2.2. Measurement of trehalose and xylose from experimental samples

Aliquots of 100 µL of fresh cucumber juice, juice extracted from cucumbers fermented in commercial vessels or from experimental media were diluted to 2 mL with water spiked with 50 µL of an internal standard of lactose (Sigma Aldrich, St. Louis, MO, USA). All solutions were filtered through OnGuard-H cartridges (Dionex Corporation, Sunnyvale, CA, USA), to remove free amino acids, into autosampler vials. The extracts were analyzed using a BioLC (Dionex Corporation) at a controlled temperature of 25 °C. The system consisted of a gradient pump, an autosampler, and a Pulsed Amperometric Detector. The mobile phase was 50 mM sodium hydroxide (NaOH) (Thermo Fisher Scientific, Fairlawn, NJ, USA) at an isocratic flow rate of 1.0 mL/min. The column used was a PA-1, 250 mm length and 4 mm i.d. (Dionex Corporation), fitted with a PA-1 Guard column (Dionex Corporation). The detector was programmed to run a quadruple waveform as recommended by the manufacturer. The detector sensitivity was set to 500 nCoulombs (nC).

The injection volume was 10 µL. Each sugar was quantified by calculating the ratio of the unknown peak height to the internal standard peak height and comparing it with a ratio of sugar standards to the internal standard (lactose). Trehalose and xylose were purchased from Fluka Chemie (Steinheim, Germany) (36).

### 2.3. Measurement of L-citrulline from experimental samples

The concentration of L-citrulline was analyzed in experimental samples using the typical procedure for free amino acids. Samples were filtered through 0.22 µ PVDF syringe filters (EMD Millipore Corp., Darmstadt, Germany). Extracts were analyzed using a Hitachi Model L-8900 Analyzer (Hitachi High Technologies, Dallas, TX, USA). The analyzer was fitted with an Ion Exchange analytical column (Hitachi # 2622SC PF, 40 mm length, 6.0 mm i.d.) connected to a guard column of the same composition. Separation of amino acids was carried out using a gradient of borate buffers (PF type, Hitachi High Technologies) and a temperature gradient of 30 to 70 °C according to the user manual supplied with the instrument with additional changes provided by Hitachi personnel (37). Post column derivatization was performed by the instrument using ninhydrin (WAKO Chemicals USA, Richmond, VA, USA). Visible detection was used at a wavelength of 570 nm. Standard curves of L-citrulline and ornithine (Sigma Aldrich, St. Louis, MO, USA) were prepared in 0.02 N hydrochloric acid (Thermo Fisher) over a range of 0.3 to 30 µg/mL. Standard curves were also prepared using an amino acid standard mix (Pierce H Standard, Thermo Pierce, Rockford, IL, USA). Ammonia was determined from the mixture. The range of concentration was 0.5 to 5 µg/mL. Under these conditions, L-citrulline eluted at 30.2 min, ornithine at 97.7 min and ammonia at 87.8 min (37).

### 2.4. Bioinformatic analysis of the genes coding for enzymes involved in trehalose, xylose and L-citrulline metabolism by certain LAB

Detection of the putative xylose symporter gene (*xylP*) and the α-glucosidase coding gene (*xylQ*) among the bacterial genomes sequenced to date was done using the Intergrated Microbial Genomics Find Gene function (38; https://img.jgi.doe.gov/cgi-bin/m/main.cgi). The analysis of the enzymes involved in the metabolism of trehalose, xylose and L-citrulline was conducted using the publicly available genome sequences for *L. pentosus* (3), *L. plantarum* (107), *L. brevis* (21), *L. buchneri* (6), and *P. pentosaceous* (7) using the Joint Genome Institute-Integrated Microbial Genomes platform (IMG; https://img.jgi.doe.gov/cgi-bin/m/main.cgi), the KEGG Orthology Pathways database (KO; http://www.kegg.jp/kegg/pathway.html) and the Metacyc (http://metacyc.org) and Biocyc (http://biocyc.org) online tools at the IMG platform as described by Ucar et al. (13).

### 2.5. Preparation of fermented cucumber juice media (FCJM)

FCJM was prepared as described by Ucar et al., 2019, *submitted*). Briefly, size 2B (32-38 mm in diameter) fresh whole pickling cucumbers from two different lots were secured from the local retail (Raleigh, NC) and fermented in a cover brine containing 80 mM calcium chloride (CaCl_2_) (Brenntag, Durham, NC, USA), 6 mM potassium sorbate (Mitsubishi International Food Ingredients, Atlanta, GA, USA), 10.1 mM calcium hydroxide (Ca(OH)_2_) (Sigma-Aldrich) and 44 mM acetic acid, added as 20% vinegar (Fleischmann Vinegar, MO, USA) to adjust the initial pH to 4.7 ± 0.1. At the end of these fermentations the pH was measured at 3.3 ± 0.1 and the media contained 0.5 ± 0.2 mM and 1.69 ± 0.6 mM glucose and fructose, respectively, as determined by HPLC conducted as described below. The cover brine and juice from this fermentation was used to prepare FCJM.

The FCJM derived from each cucumber lot were independently used and supplemented with trehalose (US Biological, cat. # T8270, 99.7% purity), xylose (Sigma-Aldrich, 99% purity) or L-citrulline (Sigma-Aldrich, 98% purity). The pH of the supplemented and un-supplemented FCJM was adjusted to 4.7 ± 0.1 or 3.7 ± 0.1 as indicated in the text using a 5N NaOH solution (Spectrum Chemicals, NJ) and 3N hydrochloric acid (HCl) (Spectrum Chemicals). pH measurements were taken using an Accumet® Research 25 pH meter (Fisher Scientific, CA, USA) equipped with a Gel-Filled Pencil-Thin pH Combination Electrode (Accumet, Fisher Scientific). The pH-adjusted FCJM were filter sterilized using 0.2-µ filtration units (Nalgene^®^-Rapid Flow^TM^, Thermo Scientific). 10 mL aliquots of each FCJM were aseptically transferred to 15 mL conical tubes for experimentation.

### 2.6. LAB cultures preparation

The bacterial cultures used for experimentation are described in Table 1. The LAB cultures were transferred from frozen stocks, prepared with Lactobacilli MRS broth (Benton, Dickinson and Company, Franklin Lakes, NJ) supplemented with 15% glycerol, to 10 mL of MRS broth. The cultures were incubated at 30 °C for 48 to 72 h prior to inoculating fermentations or the FCJM. The FCJM was inoculated to 10^5^ CFU/mL. The optical density at 600 nm of the MRS cultures was measured and used to adjust the inoculation level. A sterile 0.85% sodium chloride (NaCl) solution was used to adjust the inocula concentration as needed. Inocula with mixed cultures were prepared by combining the cells suspension in saline solution so that each strain will be at 10^5^ CFU/mL in the experimental medium.

**Table 1.** Description of the lactic acid bacteria strains used in this study.

| Genus | Species | ID Number(s) | Sources | Reference |
| --- | --- | --- | --- | --- |
| *Lactobacillus* | *plantarum* | LA0070; ATCC 14917 | Pickle cabbage | ATCC < PA Hansen < Roy Techn. Coll., Copenhagen < S. Orla-Jensen (*Streptobacterium plantarum*)<br>https://www.atcc.org/products/all/14917.aspx#history |
| *Lactobacillus* | *plantarum* | LA1196; ATCC BAA-793; NCIMB 8826; WCFS1 | Saliva | (54) |
| *Lactobacillus* | *plantarum* | 3.2.8 | Commercial cucumber fermentation | (55) |
| *Lactobacillus* | *pentosus* | LA0233; ATCC 8041 | Sauerkraut | (46) |
| *Lactobacillus* | *pentosus* | LA0445; BI0007, MOP3 | Commercial cucumber fermentation | (56) |
| *Lactobacillus* | *pentosus* | 1.8.9 | Commercial cucumber fermentation | (55) |
| *Lactobacillus* | *brevis* | LA0200; ATCC 8287 | Green Sevillano fermenting olives | RH Vaughn 269Y; (57) |
| *Lactobacillus* | *brevis* | LA0036; ATCC 14869, NRRL B-4527 | Human feces | ATCC < PA Hansen < Roy. Techn. Coll., Copenhagen < S. Orla-Jensen 14 (*Betabacterium breve*); (58) |
| *Lactobacillus* | *brevis* | 7.2.43 | Commercial cucumber fermentation | (55) |
| *Lactobacillus* | *buchneri* | LA0030; ATCC 4005, NRRL B1837 | Tomato pulp | (58) |
| *Lactobacillus* | *buchneri* | LA1149 | Commercial cucumber fermentation | (59) |
| *Lactobacillus* | *buchneri* | LA1147; E-33-07 | Commercial cucumber fermentation | (59) |

### 2.7. Analysis of the ability of certain LAB to utilize xylose, trehalose and L-citrulline in fermented cucumber juice

The fermented cucumber juice medium prepared as described above was supplemented with 18.13 ± 1.17 mM trehalose (US Biological, cat. # T8270, 99.7% purity), 18.65 ± 0.49 mM xylose (Sigma-Aldrich, cas # 58-86-6, 99% purity) or 0.56 ± 0.02 mM L-citrulline (Sigma-Aldrich, cas # 372-75-8, 98% purity) individually. FCJM supplemented with 14.21 ± 5.18 mM glucose (Sigma-Aldrich, cas # 50-99-7, 99.5% purity) or 20.61 ± 2.52 mM fructose (Sigma-Aldrich, cas # 57-48-7, 99% purity) were used as positive controls. The supplemented media were filter-sterilized using 0.2 µ filtration units (Nalgene^®^-Rapid Flow^TM^, Thermo Scientific) after adjusting the pH of the media to 4.7 ± 0.1 (Accumet® Research 25 pH meter, Fisher Scientific) using 5N NaOH (Spectrum Chemicals, NJ, USA). Aliquots of 10 mL of each supplemented FCJM were transferred to 15 mL conical plastic tubes (cat. # 430766; Corning Incorporated, Corning, NY, USA) prior to inoculation with a mixture of the three strains of *L. plantarum*, *L. pentosus*, *L. brevis* or *L. buchneri* as described in Table 1. The inocula were prepared as described above. All tubes were incubated at 30 °C for 7 days to enable the utilization of energy sources under conditions simulating post-fermentation. Negative controls were not inoculated. Samples were aseptically collected at the end point and serially diluted in 0.85% saline solution prior to plating, which was done as described below. The concentration of xylose, trehalose, and L-citrulline remaining in the media after secondary fermentation were measured by HPLC performed as described above. The concentrations of glucose, fructose, lactic acid, acetic acid and ethanol present in the FCJM after incubation for 7 days were measured using HPLC analysis as described below.

### 2.8. Enumeration of LAB from MRS agar plates

Spiral plating was conducted using an Autoplate 400 Eddy Jet 2 spiral plater (IUL, Barcelona, Spain) onto Lactobacilli deMan Rogosa and Sharpe (MRS) agar (Becton, Dickinson and Company) supplemented with 1% cycloheximide (Remel, San Diego, CA, USA) for the enumeration of LAB. MRS plates were incubated aerobically at 30 °C for 48 h. Colonies were counted using a Flash & Go Automatic Colony Counter (IUL). The limit of detection was 4 log CFU/mL.

### 2.9. Measurement of glucose, fructose, lactic acid, acetic acid and ethanol concentrations in FCJM using HPLC

The concentrations of glucose, fructose, lactic acid, acetic acid and ethanol in FCJM before and after secondary fermentation were measured using HPLC analysis. 1.5 mL of each fermentation sample was spun at 15,294 x g for 15 min at room temperature using an Eppendorf Centrifuge Model 5810 (Hamburg, Germany). A minimum of 500 µL of the supernatants were transferred into glass HPLC vials. The concentrations of the organic compounds were measured using a 30-cm HPX-87H column (Bio-Rad Laboratories, Hercules, CA, USA) and the HPLC method described by McFeeters and Barish (39) with some modifications. The operating conditions of the UFLC Shimadzu HPLC system (Shimadzu Corporation, Canby, OR, USA) included a column temperature of 65 °C and a 0.01 N sulfuric acid (H_2_SO_4_) eluent at 0.9 mL/min. A diode array detector was set at 210 nm at a rate of 1 Hz to quantify lactic acid. A RID-10A refractive index detector connected in series with the diode array detector was used to measure acetic acid, glucose, fructose and ethanol. External standardization of the detectors was done using at least five concentrations of standard compounds.

### 2.10. Evaluation of the ability of L. buchneri to utilize L-citrulline in the presence of glucose in FCJM

Fermented cucumber juice medium prepared as described above was supplemented with 12.61 ± 0.02 mM L-citrulline, 10.60 ± 0.20 mM L-citrulline and 16.49 ± 1.06 mM glucose or 12.71 ± 0.56 mM glucose and the pH adjusted to 4.7 ± 0.1. FCJM was supplemented with 10.98 ± 4.24 mM L-citrulline, 15.24 ± 0.47 mM L-citrulline and 10.83 ± 1.46 mM glucose or 10.20 ± 1.20 mM glucose in a second batch and the pH was adjusted to pH 3.7 ± 0.1. The supplemented FCJM pH was adjusted to 4.7 ± 0.1 or 3.7 ± 0.1 as described above prior to filter sterilization using 0.2 µ filtration units (Nalgene^®^-Rapid Flow™, Thermo Scientific). The filter sterilized media were aliquotted into 15 mL conical tubes (cat. # 430766; Corning Incorporated) for experimentation. Positive and negative controls were prepared without L-citrulline supplementation and non-inoculated, respectively. Aliquots of 15 mL of the experimental media were inoculated with *L. buchneri* (LA0030, LA1149, and LA1147) (Table 1). All tubes were incubated at 30 °C for 7 to 10 days, depending on the extent of growth observed. Samples for the enumeration of *L. buchneri*, pH measurement and HPLC analyses were aseptically collected and processed as described above.

### 2.11. Observation of the biochemical changes in fresh cucumber juice (FrCJ) medium inoculated with mixed starter cultures of LAB

FrCJ was prepared as described above for FCJM and mixed with cover brine in a 50:50 ratio by volume. The cover brine contained 80 mM CaCl_2_ (16.87 g), 6 mM potassium sorbate (1.71g), 44 mM acetic acid, added as 20% vinegar (Fleischmann Vinegar, MO, USA), and 0.15% Ca(OH)_2_. Two different batches of fresh cucumbers were used for the preparation of the FrCJ, that were supplemented with 18.25 ± 0.49 mM trehalose (US Biological, cat. # T8270, 99.7% purity), 51.97 ± 1.30 mM xylose (Sigma-Aldrich, cas # 58-86-6, 99% purity) and 4.23 ± 0.61 mM L-citrulline (Sigma-Aldrich, cas # 372-75-8, 98% purity). The pH of the FrCJ media was adjusted to 5.0 ± 0.1 prior to filter sterilization using 0.2 µ filtration units (Nalgene^®^-Rapid Flow™, Thermo Scientific). Aliquots of 15 mL were transferred to 50 mL conical tubes (cat. # 430829; Corning Incorporated) using aseptic techniques. The experimental tubes were inoculated with *L. pentosus* (LA0445 and 1.8.9) and *L. brevis* (3.2.19), *L. brevis* (3.2.19) alone, and a mixture of *L. pentosus* (LA0445 and 1.8.9), *L. brevis* (3.2.19), and *L. buchneri* (LA1149 and LA1147) to the levels indicated on Tables 6 and 7. The inocula were prepared as described above (Table 1). The combined inoculation volumes represented less than 10% of the FCJM volume. Un-supplemented FrCJ medium was used as a negative control and inoculated with *L. pentosus* (LA0445 and 1.8.9) and *L. brevis* (3.2.19), *L. brevis* (3.2.19) alone, and a mixture of *L. pentosus* (LA0445 and 1.8.9), *L. brevis* (3.2.19), and *L. buchneri* (LA1149 and LA1147) (Table 1). The inocula were prepared as described above. Samples were collected on days 0, 3, 7, 10, 30, 36 and 60 using aseptic techniques for spiral plating, pH measurements and HPLC analyses conducted as described above.

## 3. Results

### 3.1. Analysis of trehalose, xylose and L-citrulline content in fresh and fermented cucumbers

Concentrations of trehalose, xylose and L-citrulline were determined in fresh cucumbers and commercial cucumber fermentations. Trehalose was only detected in two out of the three samples tested collected from commercial cucumber fermentations that were 3 days old to 15.51 ± 1.68 mM (Table 2). Xylose was found in 1 out of 4 fresh cucumber samples to 36.05 mM and was not detected on samples of commercial cucumber fermentations. Decreasing concentrations of L-citrulline were found in fresh cucumbers and samples of commercial cucumber fermentations collected on days 3 and 38 to 1.65 ± 0.63, 0.85 ± 0.36 and 0.65 ± 0.07 mM, respectively (Table 2). As expected, glucose and fructose concentrations decreased as a function of the fermentation age (Table 2).

**Table 2.** Measurements of trehalose, xylose and L-citrulline in fresh and fermented cucumbers by HPLC. The limit of detection for the sugars and L-citrulline were 10 and 1.71 µM, respectively. Numbers provided in parenthesis next to the concentration values represent the number of samples analyzed in which compounds were found.

| Primary and<br>Alternative<br>Energy Sources | Concentration (mM) |  |  |
| --- | --- | --- | --- |
|  | Fresh Cucumbers<br>(n<sup>1</sup> = 4) | Day 3 of Fermentation<br>(n = 3) | Day 38 of Fermentation<br>(n = 4) |
| **Glucose** | 42.72 ± 7.4 (4) | 39.29 ± 4.07 (3) | BDL (4) |
| **Fructose** | 43.15 ± 6.5 (4) | 38.25 ± 5.13 (3) | BDL (4) |
| **Trehalose** | BDL (4) | 15.51 ± 1.68 (2) | BDL (4) |
| **Xylose** | 36.05 (1) | BDL (4) | BDL (4) |
| **L-citrulline** | 1.65 ± 0.63 (4) | 0.85 ± 0.36 (3) | 0.65 ± 0.07 (4) |

<sup>1</sup>n= number of samples analyzed

### 3.2. Bioinformatic analysis of the genes coding for enzymes involved in trehalose, xylose and L-citrulline metabolism by certain LAB

Putative *xylP* were found in *Lactobacillus antri* DSM16041, *Lactobacillus brevis* ATCC27305, *Lactobacillus hildardii* ATCC8290, *Pediococcus acidilactici* DSM 20284, two *Lactobacillus rhamnosus*, *Lactobacillus buchneri* ATCC11577 and *Lactobacillus casei* BL23. A putative *xylQ* was only found in the *Pediococcus acidilactici* genome downstream a xylose utilization operon.

Evaluation of the metabolic potential of certain LAB to utilize trehalose, xylose, and L-citrulline was done using putative metabolic pathway analysis (Table 3). Table 3 shows the results generated using 98, 3, 7, 21 and 6 genome sequences corresponding to *L. plantarum*, *L. pentosus, P. pentosaceus*, *L. brevis* and *L. buchneri*. Finished genome sequences, permanent drafts and draft sequences were used for the analysis. The metabolic potential analysis suggests that trehalose, a disaccharide, could be converted to D-glucose via the trehalose phosphorylase enzyme (EC 2.4.1.64) found in more than 97% of *L. buchneri* and *L. brevis* strains and some of the *L. plantarum* and *L. pentosus* genomes (Table 3). However, the trehalose phosphorylase putative coding gene was missing in all of the *P. pentosaceus* genomes included in the analysis (Table 3). It was also found that putatives D-xylose-5-Phosphate 3-Epimerase (EC 5.1.3.1) and Xylulokinase (EC 2.7.1.17) are commonly encoded by the *L. buchneri* genomes and less typically by the *L. plantarum* genomes (Table 3). These enzymes are involved in the conversion of D-xylulose to D-xylulose-5-phosphate via the pentose and glucuronate interconversion pathway. The genomes of several strains of *L. pentosus*, *P. pentosaceus*, and *L. brevis* were found to frequently encode for such key putative enzymes. More than 97% and 60% of the *L. plantarum* and *L. pentosus* genome sequences, respectively, were found to encode for two key enzymes in the Pentose Phosphate Pathway, the Transketolase (EC 2.2.1.1) and Transaldolase (EC 2.2.1.2), which convert D-Ribulose-5-Phosphate to D-Glyceraldehyde-3-Phosphate to feed Glycolysis; and are absent in the *L. buchneri* genome sequences (Table 3). More than 97% of the *L. plantarum*, *L. pentosus*, and *P. pentosus* genome sequences encode for enzymes involved in Glycolysis, while only 97 to 5% of the *L. brevis* genome sequences encode for the proteins involved in such pathway. The *L. buchneri* genome sequences do not encode for key enzymes involved in Glycolysis, particularly those converting D-Glyceraldehyde-3-Phosphate to smaller products.

**Table 3.**
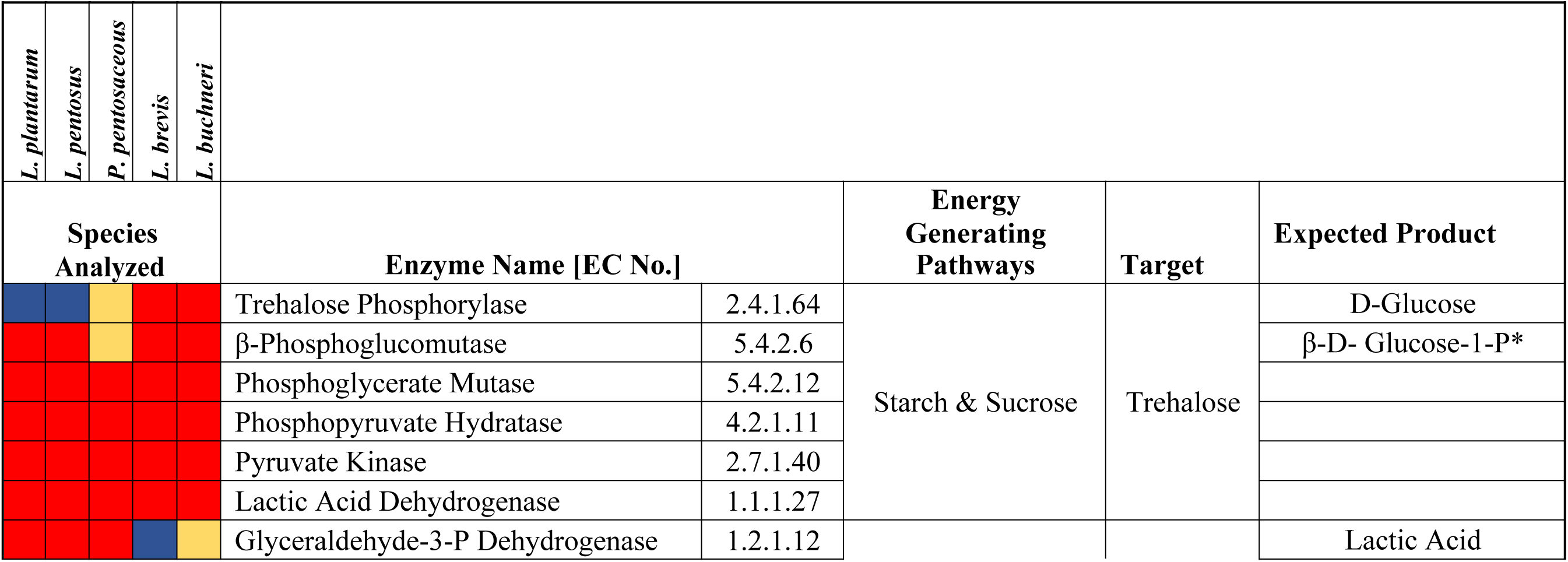
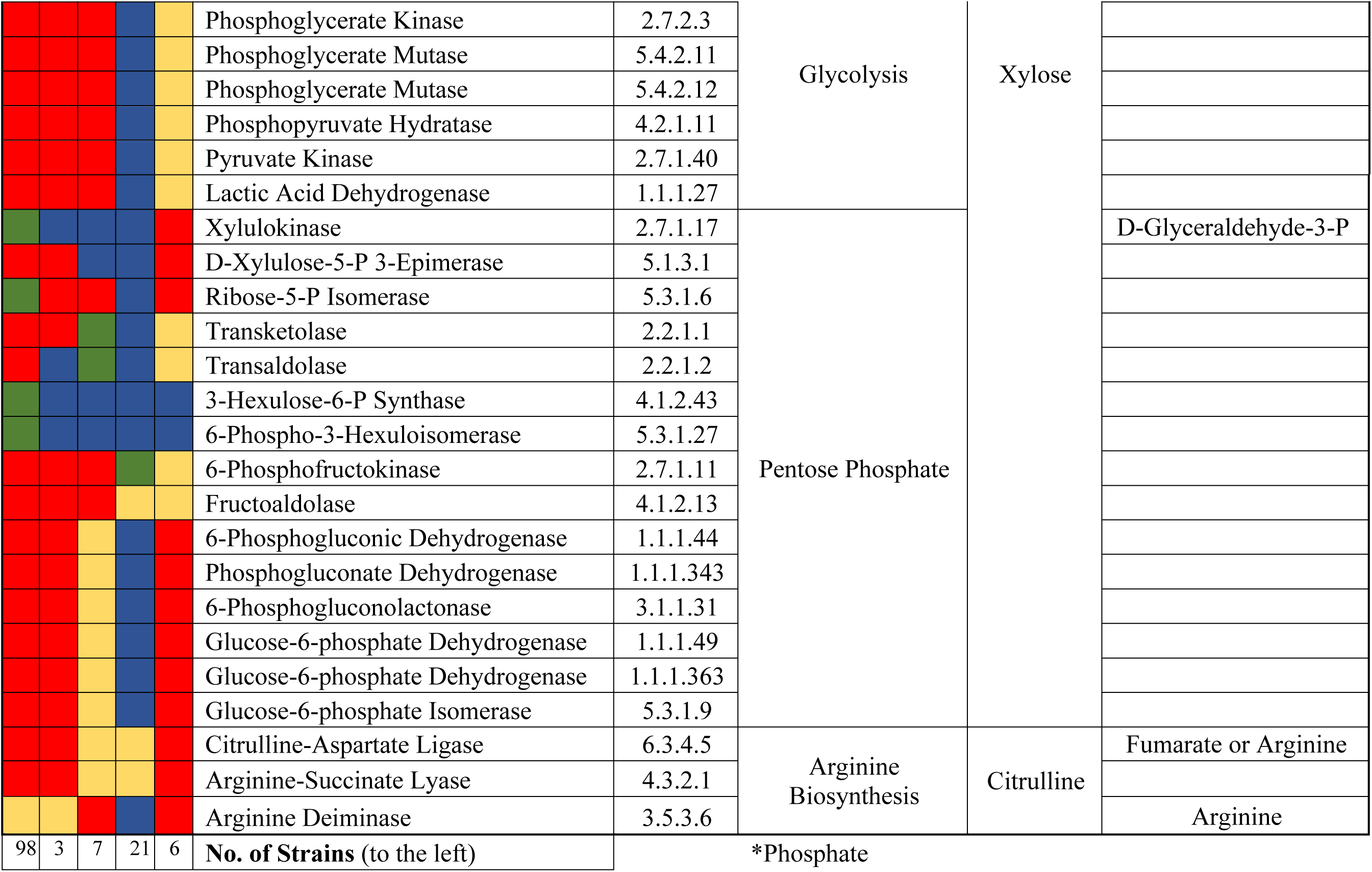
Bioinformatic analysis of the putative pathways involved in the utilization of alternative energy sources by selected lactic acid bacteria (*L. plantarum, L. pentosus, P. pentasaceus, L. brevis,* and *L. buchneri*). Each row represents the percentage of a specific lactic acid bacterium genomes that encode for the corresponding putative enzyme described on the fifth column in a metabolic pathway. All types of genome sequences were included in the analysis (finished, draft, and permanent drafts). LAB species are listed at the top of the table and the number of strains per species is listed at the bottom. The enzyme name & Enzyme Classification (EC) number, the type of pathway, target, organic compound utilized, and expected products are listed in the right four rows, respectively. The colored boxes mark the number of strains coding for a specific putative enzyme where red, blue, green and yellow represent more than 97%, between 97 and 5%, less than 5% and missing genes, respectively.

More than 97% of the *L. plantarum*, *L. pentosus*, and *L. buchneri* genome sequences included in the analysis encode for putative genes associated with L-citrulline metabolism. L-citrulline could be converted to fumarate or arginine via L-citrulline-aspartate ligase (EC 6.3.4.5) and arginine-succinate lyase (EC 4.3.2.1) (Table 3). Additionally, all the *P. pentosaceus* and *L. buchneri* genomes analyzed encode for a putative arginine deiminase which interconverts L-citrulline to arginine. More than 85% of the *L. brevis* genome sequences encode for a putative arginine deiminase (Table 3).

### 3.3. Utilization of trehalose, xylose and L-citrulline by certain LAB

The LAB that are known to homoferment or heteroferment produced mostly lactic acid or a mixture of lactic acid, acetic acid and ethanol, respectively, in FCJM supplemented with glucose or fructose (Table 4). Ethanol was not produced by the heterofermenters, *L. brevis* and *L. buchneri*, from fructose (Table 4). Substantial differences were observed in the FCJM pH and LAB colony counts by the end point of the second fermentation regardless of the species tested. However, the sum of the amounts of lactic acid, acetic acid and ethanol produced substantially surpassed the theoretical production levels based on carbon balance (Table 4). It is speculated that although the primary fermentation removed more than 99% of the detectable glucose and fructose from the system (42.72 ± 7.4 glucose and 43.15 ± 6.5 mM fructose), not all of the sugars were converted to lactic acid, acetic acid and ethanol prior to the preparation of the FCJM. An enhanced carbon balance is observed if the amounts of glucose and fructose present in the fresh cucumbers are added to those supplemented in the FCJM and total production is considered.

**Table 4.** Utilization of trehalose, xylose, and L-citrulline by selected lactic acid bacteria in fermented cucumber juice (FCJM) at pH 4.7. Trehalose, xylose and L-citrulline were supplemented to 18.13 ± 1.17, 18.65 ± 0.49 and 0.56 ± 0.02 mM, respectively. The control group glucose and fructose were supplemented to 14.21 ± 5.18 and 20.61 ± 2.52 mM, respectively. Organic compound concentrations are provided in the table below in mM. The amount of lactic acid, acetic acid and ethanol produced were calculated from the amounts measured in the FCJM after 7 days of incubation minus the amounts detected in the non-inoculated FCJM. Minimal detection limit was 10 and 1.71 µM for the sugars and L-citrulline, respectively. Ethanol was not produced from fructose, xylose or trehalose.

| Lactic Acid Bacteria Tested ➤ | *L. plantarum* | *L. pentosus* | *L. brevis* | *L. buchneri* | No Inocula<br>(Negative Control) |
| --- | --- | --- | --- | --- | --- |
| **Glucose** |  |  |  |  |  |
| Glucose Utilized | $14.71 \pm 5.38^a$ | $14.24 \pm 5.22^a$ | $12.66 \pm 4.50^b$ | $14.71 \pm 5.38^a$ | NA<sup>*</sup> |
| Lactic Acid Produced | $79.01 \pm 6.51^a$ | $88.43 \pm 6.62^b$ | $43.05 \pm 9.45^c$ | $33.39 \pm 0.97^d$ | $103.02 \pm 3.71$ |
| Acetic Acid Produced | BDL<sup>a</sup> | BDL<sup>a</sup> | $7.06 \pm 2.31^b$ | $7.54 \pm 0.25^b$ | $35.90 \pm 0.96$ |
| Ethanol Produced | BDL<sup>a</sup> | BDL<sup>a</sup> | $8.33 \pm 0.03^b$ | $9.10 \pm 2.07^b$ | $9.73 \pm 0.03$ |
| Final pH | $3.91 \pm 0.00^a$ | $3.89 \pm 0.01^a$ | $4.32 \pm 0.04^b$ | $4.24 \pm 0.04^b$ | $4.55 \pm 0.03$ |
| Growth (Log CFU/mL) | $\geq 8$ | $\geq 8$ | $\geq 8$ | $8.83 \pm 0.20$ | NA |
| **Fructose** |  |  |  |  |  |
| Fructose Utilized | $22.30 \pm 3.12^a$ | $22.30 \pm 3.12^a$ | $21.36 \pm 2.74^a$ | $21.98 \pm 2.98^a$ | NA |
| Lactic Acid Produced | $70.34 \pm 9.38^a$ | $67.90 \pm 5.83^a$ | $35.58 \pm 13.01^b$ | $5.06 \pm 1.30^c$ | $99.34 \pm 0.55$ |
| Acetic Acid Produced | BDL<sup>a</sup> | BDL<sup>a</sup> | $9.49 \pm 0.41^b$ | $8.03 \pm 3.64^b$ | $35.41 \pm 0.00$ |
| Final pH | $3.9 \pm 0.05^a$ | $3.89 \pm 0.04^a$ | $4.35 \pm 0.04^b$ | $4.58 \pm 0.04^b$ | $4.55 \pm 0.03$ |
| Growth (Log CFU/mL) | $\geq 8$ | $\geq 8$ | $\geq 8$ | $8.78 \pm 0.29$ | NA |
| **Xylose** |  |  |  |  |  |
| Xylose Utilized | BDL<sup>a</sup> | $3.14 \pm 0.37^b$ | $16.92 \pm 0.23^c$ | $18.65 \pm 0.49^c$ | NA |
| Lactic Acid Produced | BDL<sup>a</sup> | $29.54 \pm 18.70^b$ | $30.77 \pm 4.85^b$ | $9.74 \pm 12.16^c$ | $98.96 \pm 1.20$ |
| Acetic Acid Produced | BDL<sup>a</sup> | BDL<sup>a</sup> | $6.83 \pm 4.42^b$ | $13.51 \pm 2.18^c$ | $39.99 \pm 0.01$ |
| Final pH | $4.26 \pm 0.01^a$ | $4.21 \pm 0.01^a$ | $4.19 \pm 0.04^a$ | $4.24 \pm 0.05^a$ | $4.55 \pm 0.03$ |
| Growth (Log CFU/mL) | $< 4$ | $< 4$ | $\geq 8$ | $8.53 \pm 0.02$ | NA |

| Trehalose |  |  |  |  |  |
| --- | --- | --- | --- | --- | --- |
| Trehalose Utilized | $18.13 \pm 1.17^a$ | $13.11 \pm 1.04^b$ | BDL<sup>c</sup> | $2.36 \pm 1.44^d$ | NA |
| Lactic Acid Produced | $94.05 \pm 3.14^a$ | $63.28 \pm 22.02^b$ | BDL<sup>c</sup> | $3.72 \pm 5.22^d$ | $98.96 \pm 0.01$ |
| Acetic Acid Produced | BDL<sup>a</sup> | BDL<sup>a</sup> | BDL<sup>a</sup> | $10.99 \pm 4.77^b$ | $22.09 \pm 0.56$ |
| Final pH | $3.7 \pm 0.07^a$ | $3.87 \pm 0.06^a$ | $4.5 \pm 0.01^b$ | $4.56 \pm 0.01^b$ | $4.55 \pm 0.03$ |
| Growth (Log CFU/mL) | $\geq 8$ | $\geq 8$ | $< 4$ | $8.47 \pm 0.03$ | NA |
| L-citrulline |  |  |  |  |  |
| Remaining Citrulline | $0.73 \pm 0.01^a$ | $0.72 \pm 0.04^a$ | $0.76 \pm 0.18^a$ | $0.08 \pm 0.01^b$ | $0.59 \pm 0.07$ |
| Final pH | $4.3 \pm 0.01^a$ | $4.31 \pm 0.01^a$ | $4.15 \pm 0.19^b$ | $4.49 \pm 0.02^c$ | $4.51 \pm 0.01$ |
| Growth (Log CFU/mL) | $\geq 8$ | $\geq 8$ | $\geq 8$ | $8.39 \pm 0.08$ | NA |

\* Not Applicable
\*\* The results in the rows within blocks followed by the same alphabets (a, b, c, and d) indicate that they are not significantly different ( $p > 0.05$ ) from each other and the treatments with the different alphabets indicate that they are significantly different ( $p < 0.05$ ).
<sup>‡</sup>BDL: Below Detection Level

Thus, approximately 57 mM glucose and 43 mM fructose were converted to about 182 and 191 mM lactic acid by *L. plantarum* and *L. pentosus*, respectively, in the FCJM supplemented with glucose and 43 mM glucose and 63 mM fructose were converted to 169 and 167 mM lactic acid by *L. plantarum* and *L. pentosus*, respectively, in the FCJM supplemented with fructose. A similar carbon balance is observed for the other LAB tested if acetic acid and ethanol production is also considered. About 20 to 30 mM of the hexose-derived carbon was still not accounted for, which was presumed to be assimilated by the biomass.

*L. brevis* (1.73±0.72 mM) and *L. buchneri* (<0.01 mM) utilized nearly all the xylose supplemented in the FCJM (18.65 ± 0.49) while *L. plantarum* and *L. pentosus* did not (Table 4). *L. brevis* and *L. buchneri* converted xylose to lactic acid and acetic acid but not ethanol, dropped the pH from 4.7 ± 0.1 to about 4.2 and doubled their cell densities, reaching colony counts above 8 log CFU/mL (Table 4). On the contrary, no increases in cell densities were observed in FCJM inoculated with *L. plantarum* or *L. pentosus*, suggesting such cells were not energized by xylose (Table 4).

The opposite pattern from xylose utilization was observed for trehalose catabolism. Trehalose was converted to lactic acid by *L. plantarum* and *L. pentosus* with the consequent decrease in pH to 3.7 and an increase in cell density above 8 log CFU/mL (Table 4). *L. brevis* did not utilize trehalose and *L. buchneri* partially utilized trehalose (2.36 ± 1.44) converting it to lactic and acetic acids and deriving sufficient energy for growth and some acidification (Table 4).

*L. buchneri* was unique in utilizing L-citrulline and deriving energy for growth with minimal changes in pH (Table 4). An increase in the FCJM L-citrulline concentration was measured from samples inoculated with *L. plantarum*, *L. pentosus* and *L. brevis* with significant decreases in pH and increases in colony counts (Table 4). Although, *L. plantarum*, *L. pentosus* and *L. brevis* were unable to utilize L-citrulline, they produced small of amounts of the amino acid and were able to grow in the FCJM supplemented with it (Table 4).

### 3.4. Analysis of the ability of L. buchneri to utilize L-citrulline in the presence of excess and limiting glucose

Utilization of L-citrulline by *L. buchneri* in FCJM with an adjusted pH of 4.7 ± 0.1 and supplemented with 12.61 ± 0.02 mM of the amino acid resulted in an increase in colony counts and pH from 4.7 ± 0.1 to 5.18 ± 0.01 and the production of acetic acid, ornithine and ammonia (Table 5). A slight decrease in pH was observed when L-citrulline was co-utilized with glucose by *L. buchneri* in FCJM at pH 4.7, supplemented with the amino acid and the sugar. Co-metabolism of 10.44 ± 0.9 mM L-citrulline with 16.49 ± 5.3 mM glucose prevented the formation of ornithine and ammonia and resulted in the equimolar production of lactic acid (21.41 ± 9.84 mM) and acetic acid (20.72 ± 12.08 mM) (Table 5). Production of ammonia and ornithine was observed in the non-supplemented FCJM in which an increase in colony counts of 2 log of CFU/mL was detected (Table 5). Supplementation of the FCJM with 16.49 ± 1.06 mM glucose resulted in some production of acetic and lactic acids within the 10 days of incubation with minimal changes in pH and colony counts (Table 5).

**Table 5.**
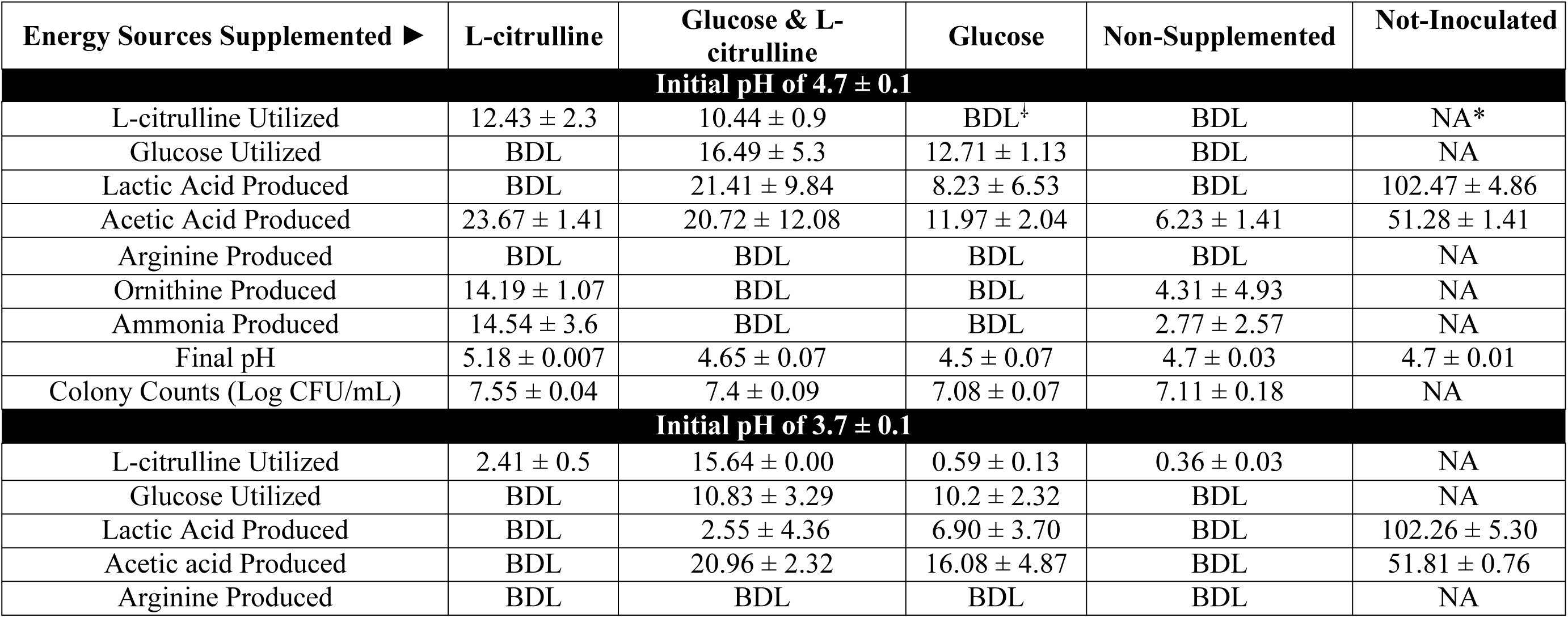
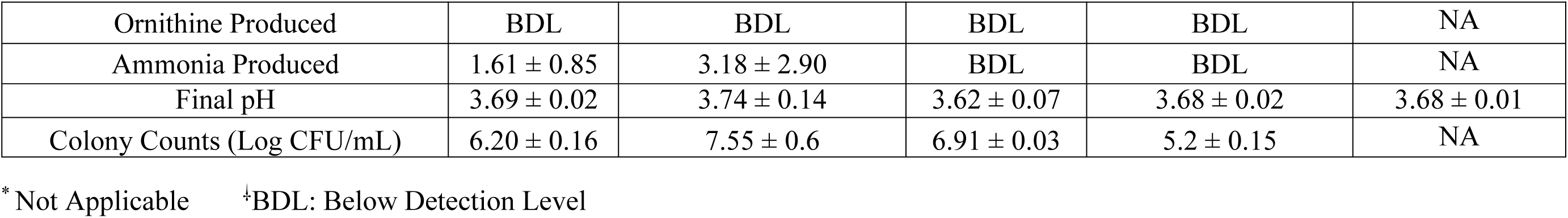
Utilization of L-citrulline by *Lactobacillus buchneri* in the presence of glucose. The initial fermented cucumber juice medium pH was adjusted to 4.7 or 3.7 ± 0.1. Values for metabolic end products (mM), pH and colony counts determined from Lactobacilli MRS Agar plates are shown. The amount of lactic acid and acetic acid produced were calculated from the total amounts easured in the FCJM from samples collected after seven days of incubation minus the concentrations detected in the Non-Inoculated FCJM. While no significant difference was determined between the treatments and control values for cultures with an initial pH of 3.7 ± 0.1 (p>0.05), a significant difference was observed in the pH values measured from treatments and control corresponding to cultures with an initial pH of 4.7 ± 0.1 (p<0.05) using an ANOVA test. Minimal limit of detection for metabolites was < 10 µM.

| Energy Sources Supplemented ► | L-citrulline | Glucose & L-citrulline | Glucose | Non-Supplemented | Not-Inoculated |
| --- | --- | --- | --- | --- | --- |
| **Initial pH of $4.7 \pm 0.1$** |  |  |  |  |  |
| L-citrulline Utilized | $12.43 \pm 2.3$ | $10.44 \pm 0.9$ | BDL<sup>†</sup> | BDL | NA* |
| Glucose Utilized | BDL | $16.49 \pm 5.3$ | $12.71 \pm 1.13$ | BDL | NA |
| Lactic Acid Produced | BDL | $21.41 \pm 9.84$ | $8.23 \pm 6.53$ | BDL | $102.47 \pm 4.86$ |
| Acetic Acid Produced | $23.67 \pm 1.41$ | $20.72 \pm 12.08$ | $11.97 \pm 2.04$ | $6.23 \pm 1.41$ | $51.28 \pm 1.41$ |
| Arginine Produced | BDL | BDL | BDL | BDL | NA |
| Ornithine Produced | $14.19 \pm 1.07$ | BDL | BDL | $4.31 \pm 4.93$ | NA |
| Ammonia Produced | $14.54 \pm 3.6$ | BDL | BDL | $2.77 \pm 2.57$ | NA |
| Final pH | $5.18 \pm 0.007$ | $4.65 \pm 0.07$ | $4.5 \pm 0.07$ | $4.7 \pm 0.03$ | $4.7 \pm 0.01$ |
| Colony Counts (Log CFU/mL) | $7.55 \pm 0.04$ | $7.4 \pm 0.09$ | $7.08 \pm 0.07$ | $7.11 \pm 0.18$ | NA |
| **Initial pH of $3.7 \pm 0.1$** |  |  |  |  |  |
| L-citrulline Utilized | $2.41 \pm 0.5$ | $15.64 \pm 0.00$ | $0.59 \pm 0.13$ | $0.36 \pm 0.03$ | NA |
| Glucose Utilized | BDL | $10.83 \pm 3.29$ | $10.2 \pm 2.32$ | BDL | NA |
| Lactic Acid Produced | BDL | $2.55 \pm 4.36$ | $6.90 \pm 3.70$ | BDL | $102.26 \pm 5.30$ |
| Acetic acid Produced | BDL | $20.96 \pm 2.32$ | $16.08 \pm 4.87$ | BDL | $51.81 \pm 0.76$ |
| Arginine Produced | BDL | BDL | BDL | BDL | NA |
| Ornithine Produced | BDL | BDL | BDL | BDL | NA |
| Ammonia Produced | $1.61 \pm 0.85$ | $3.18 \pm 2.90$ | BDL | BDL | NA |
| Final pH | $3.69 \pm 0.02$ | $3.74 \pm 0.14$ | $3.62 \pm 0.07$ | $3.68 \pm 0.02$ | $3.68 \pm 0.01$ |
| Colony Counts (Log CFU/mL) | $6.20 \pm 0.16$ | $7.55 \pm 0.6$ | $6.91 \pm 0.03$ | $5.2 \pm 0.15$ | NA |

\* Not Applicable ‡BDL: Below Detection Level

**Table 6.**
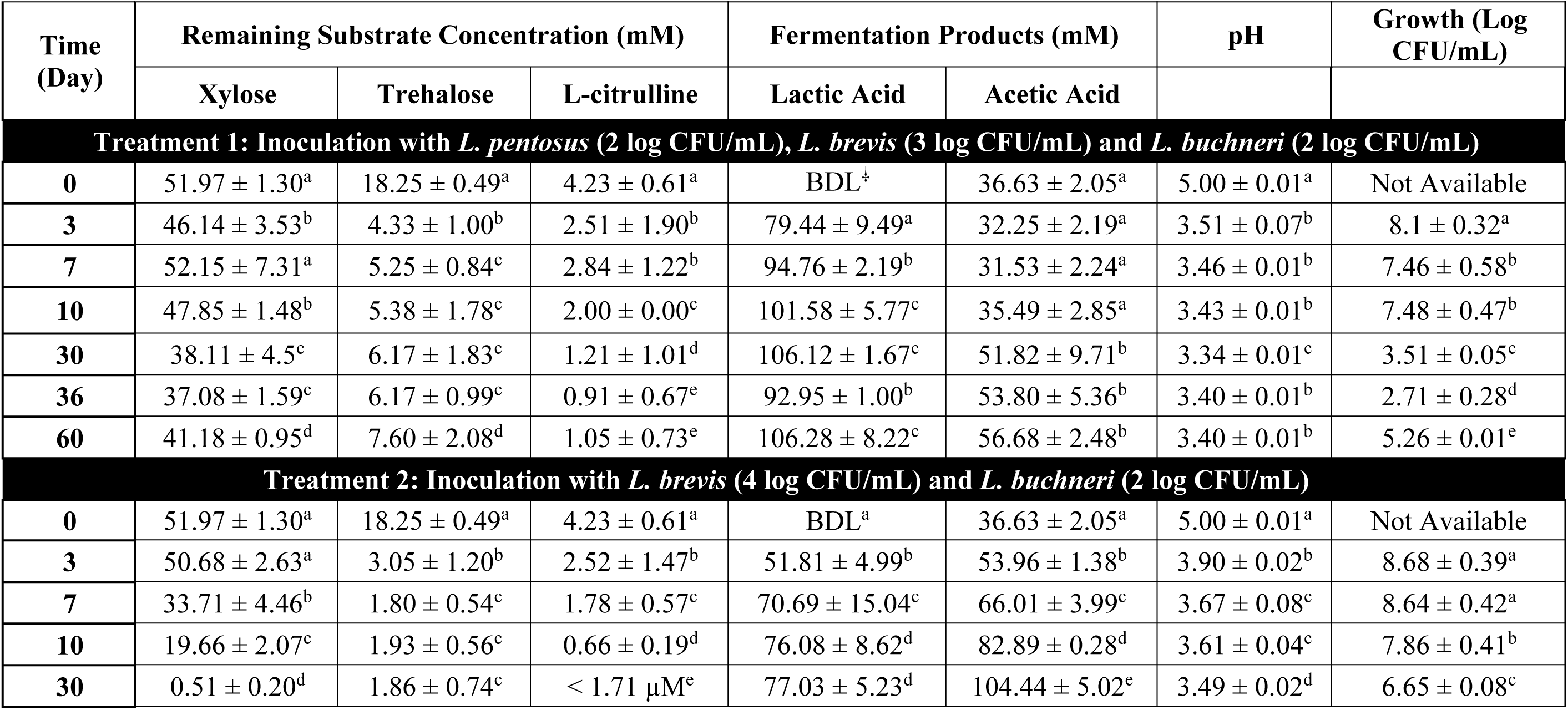
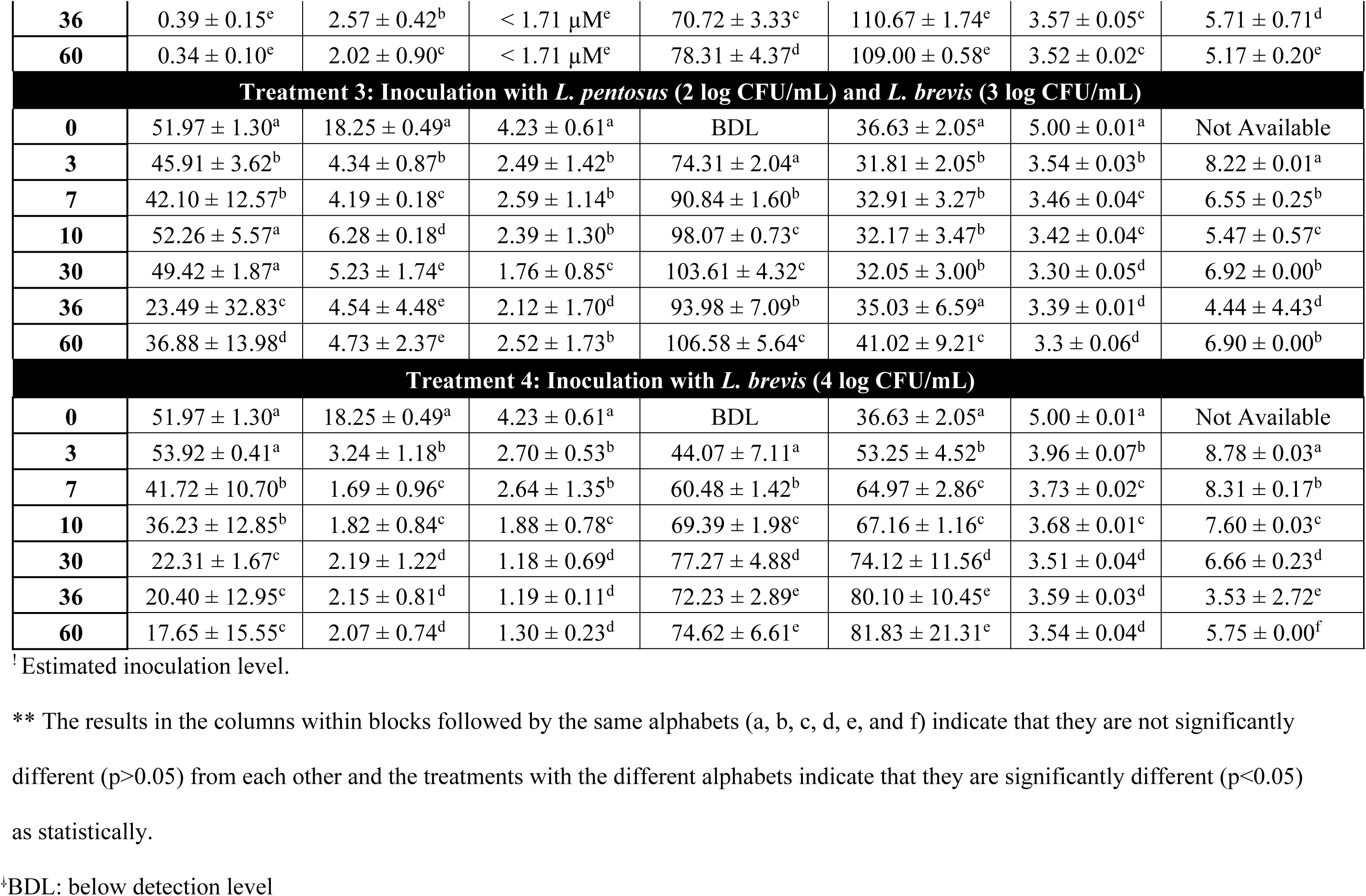
Fermentation by certain LAB of fresh cucumber juice supplemented with 18.25 ± 0.49 mM trehalose, 51.97 ± 1.30 mM xylose and 4.23 ± 0.61 mM L-citrulline. The starter cultures were inoculated to variable levels. *L. pentosus* (LA0455 and 1.8.9), *L. brevis* (3.2.19) and *L. buchneri* (LA1149 and LA1147) were used for inoculation. The FCJ medium initial pH was 5.0 ± 0.1. Minimal detection limits for the fermentation metabolites using HPLC was 0.01 mM. There were 10.30 ± 3.56 mM glucose and 12.16 ± 5.35 mM fructose present in this FrCJ medium derived from the fresh cucumber juice used to prepare the culture medium.

**Table 7.** Fermentation by certain LAB of raw fresh cucumber juice medium inoculated with mixed starter cultures: The starter cultures were inoculated to variable levels. *L. pentosus* (LA0455 and 1.8.9), *L. brevis* (3.2.19) and *L. buchneri* (LA1149 and LA1147) were used for inoculation. The FCJ medium initial pH was 5.0 ± 0.1. There were 10.30 ± 3.56 mM glucose and 12.16 ± 5.35 mM fructose present in this FrCJ medium. ^!^ Estimated inoculation level. ^⸸^ BDL: below detection level ** The results in the columns within blocks followed by the same alphabets (a, b, c, d, e, and f) indicate that they are not significantly different (p>0.05) from each other and the treatments with the different alphabets indicate that they are significantly different (p<0.05) as statistically.

| ime<br>(day) | Remaining Concentration (mM) |  |  | Products Concentration (mM) |  | pH | Growth (log<br>CFU/mL) |
| --- | --- | --- | --- | --- | --- | --- | --- |
|  | Xylose | Trehalose | L-citrulline | Lactic Acid | Acetic Acid |  |  |
| Treatment 1: Inoculation with *L. pentosus* (2 log CFU/mL) and *L. brevis* (3 log CFU/mL) |  |  |  |  |  |  |  |
| 0 | < 0.01 | < 0.01 | 0.39 ± 0.11<sup>a</sup> | BDL<sup>‡</sup> | 34.01 ± 2.83<sup>a</sup> | 5.00 ± 0.01<sup>a</sup> | Not Available |
| 3 | 1.90 ± 1.14 | 0.08 ± 0.04 | 0.38 ± 0.18<sup>a</sup> | 59.66 ± 4.01<sup>a</sup> | 33.65 ± 2.23<sup>a</sup> | 3.60 ± 0.01<sup>b</sup> | 8.03 ± 0.49<sup>a</sup> |
| 7 | < 0.01 | < 0.01 | 0.56 ± 0.10<sup>b</sup> | 62.45 ± 0.08<sup>a</sup> | 33.77 ± 2.32<sup>a</sup> | 3.59 ± 0.03<sup>b</sup> | 7.24 ± 0.00<sup>b</sup> |
| 10 | < 0.01 | < 0.01 | 0.45 ± 0.09<sup>c</sup> | 64.23 ± 4.30<sup>a</sup> | 33.67 ± 3.24<sup>a</sup> | 3.60 ± 0.02<sup>b</sup> | 5.28 ± 0.08<sup>c</sup> |
| 30 | < 0.01 | < 0.01 | 0.41 ± 0.15<sup>c</sup> | 65.82 ± 0.61<sup>a</sup> | 32.08 ± 3.13<sup>a</sup> | 3.48 ± 0.04<sup>b</sup> | 5.71 ± 0.78<sup>d</sup> |
| 36 | < 0.01 | < 0.01 | 0.38 ± 0.03<sup>a</sup> | 65.00 ± 7.04<sup>a</sup> | 33.43 ± 1.50<sup>a</sup> | 3.52 ± 0.01<sup>b</sup> | 4.11 ± 1.48<sup>e</sup> |
| 60 | 0.05 ± 0.01 | < 0.01 | 0.52 ± 0.29<sup>b</sup> | 65.66 ± 5.73<sup>a</sup> | 33.73 ± 4.22<sup>a</sup> | 3.51 ± 0.06<sup>b</sup> | 4.44 ± 0.40<sup>f</sup> |
| Treatment 2: Inoculation with *L. brevis* (4 log CFU/mL) |  |  |  |  |  |  |  |
| 0 | < 0.01 | < 0.01 | 0.39 ± 0.11<sup>a</sup> | BDL | 34.01 ± 2.83<sup>a</sup> | 5.00 ± 0.01<sup>a</sup> | Not Available |
| 3 | 2.33 ± 0.31 | < 0.01 | 0.35 ± 0.18<sup>a</sup> | 24.37 ± 3.49<sup>a</sup> | 38.78 ± 1.16<sup>b</sup> | 4.26 ± 0.01<sup>b</sup> | 8.42 ± 0.18<sup>a</sup> |
| **7** | 0.54 ± 0.07 | < 0.01 | 0.33 ± 0.11<sup>a</sup> | 25.84 ± 2.34<sup>a</sup> | 39.32 ± 2.76<sup>b</sup> | 4.33 ± 0.00<sup>c</sup> | 7.23 ± 0.03<sup>b</sup> |
| **10** | < 0.01 | < 0.01 | 0.34 ± 0.07<sup>a</sup> | 26.65 ± 3.01<sup>a</sup> | 38.20 ± 3.69<sup>b</sup> | 4.25 ± 0.06<sup>b</sup> | 7.00 ± 0.09<sup>b</sup> |
| **30** | < 0.01 | < 0.01 | < 1.71 µM | 37.98 ± 20.02<sup>b</sup> | 40.63 ± 0.93<sup>b</sup> | 3.97 ± 0.18<sup>d</sup> | 5.99 ± 0.82<sup>c</sup> |
| **36** | < 0.01 | < 0.01 | < 1.71 µM | 37.52 ± 15.64<sup>b</sup> | 41.76 ± 2.60<sup>b</sup> | 4.11 ± 0.28<sup>e</sup> | 5.11 ± 0.17<sup>d</sup> |
| **60** | < 0.01 | < 0.01 | < 1.71 µM | 34.00 ± 17.33<sup>b</sup> | 37.06 ± 1.30<sup>b</sup> | 4.06 ± 0.18<sup>e</sup> | 4.71 ± 0.02<sup>e</sup> |
| **Treatment 3: Inoculation with *L. pentosus* (2 log CFU/mL), *L. brevis* (3 log CFU/mL) and *L. buchneri* (2 log CFU/mL)** |  |  |  |  |  |  |  |
| **0** | < 0.01 | < 0.01 | 0.39 ± 0.11<sup>a</sup> | BDL | 34.01 ± 2.83<sup>a</sup> | 5.00 ± 0.01<sup>a</sup> | Not Available |
| **3** | 2.73 ± 1.38 | 0.09 ± 0.05 | 0.44 ± 0.07<sup>b</sup> | 58.16 ± 1.76<sup>a</sup> | 33.02 ± 1.58<sup>a</sup> | 3.63 ± 0.03<sup>b</sup> | 7.91 ± 0.40<sup>a</sup> |
| **7** | < 0.01 | < 0.01 | 0.48 ± 0.09<sup>b</sup> | 64.02 ± 1.79<sup>b</sup> | 32.90 ± 0.90<sup>a</sup> | 3.61 ± 0.04<sup>b</sup> | 6.68 ± 0.10<sup>b</sup> |
| **10** | 0.04 ± 0.01 | < 0.01 | 0.33 ± 0.18<sup>a</sup> | 63.32 ± 6.36<sup>b</sup> | 33.25 ± 2.79<sup>a</sup> | 3.61 ± 0.01<sup>b</sup> | 7.06 ± 0.09<sup>c</sup> |
| **30** | < 0.01 | < 0.01 | < 1.71 µM | 59.78 ± 3.50<sup>a</sup> | 33.32 ± 3.96<sup>a</sup> | 3.57 ± 0.01<sup>b</sup> | 5.48 ± 0.18<sup>d</sup> |
| **36** | < 0.01 | < 0.01 | < 1.71 µM | 56.80 ± 7.31<sup>a</sup> | 36.03 ± 3.07<sup>a</sup> | 3.57 ± 0.01<sup>b</sup> | 5.26 ± 0.04<sup>e</sup> |
| **60** | < 0.01 | < 0.01 | < 1.71 µM | 58.65 ± 6.63<sup>a</sup> | 36.05 ± 3.48<sup>a</sup> | 3.64 ± 0.01<sup>b</sup> | 5.02 ± 0.24<sup>f</sup> |

Minimal changes in the concentration of L-citrulline, pH and ammonia production were observed after 10 days of incubation in the FCJM at pH 3.7, supplemented with L-citrulline and inoculated with *L. buchneri* (Table 5). However, there was a 1 log of CFU/mL increase in colony counts from MRS agar plates. Supplementation of the FCJM with glucose and L-citrulline (pH 3.7 ± 0.1) enabled a 2.5 log of CFU/mL increase in colony counts, production of lactic acid (2.55 ± 4.36), acetic acid (20.96 ± 2.32) and ammonia (3.18 ± 2.90) and an ending pH of 3.74 ± 0.14 (Table 5). The absence of L-citrulline in FCJM supplemented with glucose (pH 3.7 ± 0.1) resulted in the utilization of 12.71 ± 1.13 mM of the hexose, which was converted to 6.90 ± 3.70 mM lactic acid and 11.97 ± 2.04 mM acetic acid with an ending pH of 3.62 ± 0.07 after 7 days of incubation (Table 5). In the absence of supplements in the FCJM at pH 3.7, *L. buchneri* did not proliferate, maintaining a colony count at 5.2 ± 0.12 (Table 5).

### 3.5. Observation of the biochemical changes in fresh cucumber juice (FrCJ) medium supplemented with xylose, trehalose and L-citrulline and inoculated with mixed starter cultures of LAB

Treatments 1, 2 and 3 were co-inoculated with *L. pentosus* LA0445 and 1.8.9, *L. brevis* 3.2.19 and *L. buchneri* LA1147 and LA1149. The *L. pentosus* and *L. brevis* strains were inoculated to 2, 3 and 4 log CFU/mL and 3, 4 and 5 log CFU/mL, respectively, in the FrCJ medium. Essentially, *L. pentosus* was inoculated 1 log CFU/mL below the inoculation level for *L. brevis*, given its shown robustness in cucumber fermentations (40). *L. buchneri* strains, a spoilage associated LAB for fermented cucumbers, were inoculated to 2 log CFU/mL in all treatments. No substantial differences in substrate utilization trends or fermentation end products were observed as a function of *L. pentosus* or *L. brevis* inoculation level and thus only data for treatment 1 is shown in Table 6. Utilization of trehalose, L-citrulline and xylose was observed in treatments 1, 2 and 3 with the disaccharide as the preferred substrate over xylose, but not L-citrulline (Table 6). The FrCJ medium pH decreased from 5.00 ± 0.01 to 3.40 ± 0.01 after 36 days of incubation (Table 6). Colony counts from MRS agar plates increased from 5.0 log CFU/mL to 8.1 ± 0.32 log CFU/mL by day 3 and were at 7.48 ± 0.47 log CFU/mL on day 10 (Table 6). Cell densities decreased to 3.51 ± 0.05 and 2.71 ± 0.28 log CFU/mL by days 30 and 36, respectively. However, a second increment in colony counts from MRS plates was observed on day 60 to 5.26 ± 0.01 log CFU/mL in the FrCJ (Table 6). The exclusion of *L. pentosus* from the inocula in treatment 4 resulted in a more complete utilization of the alternate energy sources, xylose, trehalose and L-citrulline after 10 days of incubation, an end fermentation pH of 3.5 ± 0.02, slightly higher than the standard end of fermentation pH which fluctuates between 3.3 and 3.0 and the highest production of acetic and lactic acids at about 157 mM as compared to 117 to 125 mM in the other treatments (Table 6). Xylose utilization was initiated earlier in the fermentation, 27.97 ± 3.27 mM less lactic acid was produced, the amount of acetic acid formed doubled as compared to treatment 1 and a steady drop in colony counts to 5.17 ± 0.20 occurred by 60 days of incubation (Table 6). Exclusion of *L. buchneri* from the inocula used in treatment 1, essentially inoculation with *L. pentosus* and *L. brevis* presented trends in substrate utilization and fermentation end product formation that were similar to treatment 1, except for a reduction in acetic acid production by about 15 mM and some fluctuations in colony counts after 30 days of incubation (Table 6). Treatment 6 was inoculated with *L. brevis* alone to 4 log CFU/mL. *L. brevis* utilized more xylose than the corresponding amount utilized in treatment 1 but less than in treatment 4, which also contained *L. buchneri* (Table 6). L-citrulline was also removed by *L. brevis* in treatment 6, which presented a final pH of 3.54 ± 0.04 (Table 6).

### 3.6. Observation of the biochemical changes in unsupplemented FrCJ medium inoculated with mixed starter cultures of LAB

As suggested by the data presented in Table 2, only L-citrulline was detected in the FrCJ medium utilized for this experiment (Table 7). Inoculation of FrCJ medium with *L. pentosus* to 2 log CFU/mL and *L. brevis* to 3 log CFU/mL (treatment 1) resulted in the presence of traces of L-citrulline (0.52 ± 0.29 mM) in the medium after 60 days of incubation, production of 65.66 ± 5.73 mM lactic acid and 33.73 ± 4.22 mM acetic acid and an ending pH of 3.51 ± 0.06 with viable colony counts from MRS plates at 4.44 ± 0.40 log CFU/mL (Table 7). Inoculation of the FrCJ medium with *L. brevis* alone to 4 log CFU/mL (treatment 2) resulted in an incomplete fermentation with the production of 34.00 ± 17.33 mM lactic acid, 37.06 ± 1.30 mM acetic acid, an ending pH of 4.06 ± 0.18 and viable colony counts from MRS agar plates at 4.71 ± 0.02 (Table 7). However, use of a tripartite starter culture of *L. pentosus* inoculated to 2 log of CFU/mL, *L. brevis* inoculated to 3 log CFU/mL and *L. buchneri* inoculated to 2 log CFU/mL (treatment 3) resulted in the complete removal of L-citrulline, mostly occurring between days 10 and 30, and the production of 58.65 ± 6.63 mM lactic acid and 36.05 ± 3.48 mM acetic acid with an end of fermentation pH around 3.64 ± 0.01 and viable colony counts at 5.02 ± 0.24 log CFU/mL (Table 7). No changes in lactic and acetic acids concentrations were observed in the FrCJ medium inoculated with the three cultures after day 10 (Table 7). Colony counts for treatment 3 did not increase after day 10 either (Table 7).

## 4. Discussion

This study investigated the utilization of alternative energy sources by the organisms prevailing in cucumber fermentations, including *L. pentosus*, *L. plantarum*, *L. brevis* and *P. pentosaceous*. It was hypothesized that utilization of alternate energy sources by the LAB prevailing in cucumber fermentation would hamper the ability of spoilage associated microbes such as *L. buchneri* to derive energy for growth and/or metabolic activity post-fermentation. The amounts of potential alternate energy sources in commercial cucumber fermentations were determined. It was found that xylose is occasionally present in fresh cucumbers but disappears from fermentations before day 3 (Table 2). Trehalose is often produced between day 1 and 3 of commercial cucumber fermentations, suggesting that the indigenous microbiota is responding to the osmotic stress in cover brines containing at least 5.8 % sodium chloride (NaCl) in an effort to retain viability (14, 41, 16, 15). L-citrulline is present in fresh cucumbers and at the end of commercial fermentations (Table 2). Thus, the removal of L-citrulline becomes a target in the prevention of growth of spoilage organisms such as *L. buchneri* during or after fermentation.

To define the ability of certain LAB to utilize potential alternate energy sources a FCJM was used as a model system to mimic conditions post-fermentation. The LAB tested were able to metabolize glucose and fructose in FCJM and generate an increase in cell densities of about 2.5 to 3 log CFU/mL (Table 4). This observation suggests there are sufficient essential nutrients for growth of LAB in FCJM and that the reduction in pH after a fermentation is apparently completed is the main factor stopping further proliferation and lactic acid production. The lack of carbon balance using the data collected from the fermentation of FCJM supplemented with glucose and fructose suggest an uncoupling with regards to the consumption of the sugars and its conversion to end products such as lactic acid, acetic acid and ethanol (Table 4). Such uncoupling highlights the need to consider a cucumber fermentation completed only after all the carbon is accounted for instead of the time when glucose and fructose are no longer detectable.

The same imbalance in the carbon consumed vs. produced applied towards the end of the FCJM fermentation, which was still missing between 20 and 30 mM of the carbon consumed as hexoses (Table 4). Nevertheless, the fact that LAB can proliferate in FCJM if the pH is adjusted above 3.3 ± 0.1, suggest that fermented cucumbers are more microbiologically unstable than it was presumed.

As expected, trehalose was utilized in FCJM under conditions similar to those present after cucumber fermentations are completed by the LAB that prevail in commercial cucumber fermentations, *L. plantarum* and *L. pentosus* (40), and to a lesser extent by *L. buchneri*. Trehalose uptake by lactobacilli and pediococci is facilitated by phosphotransferases and intracellular phosphor-glycosyl hydrolases (42, 21, 43, 41, 26, 44, 45, 6). The putative metabolic potential to utilize trehalose was identified in this study for 97% of the *L. buchneri* and *L. brevis* genome sequences included in the bioinformatic analysis and some of the *L. plantarum* and *L. pentosus* genome sequences, but not for the *P. pentosaceus* genome sequences. In line with the observations made by others, *L. brevis* did not utilize trehalose in FCJM even though the putative enzymes involved in its metabolism were found in the genome sequences studied (Table 3) (21, 26, 44, 45, 6). The *L. brevis* utilized in this study were isolated from vegetable fermentations and beer and are underrepresented in the pool of genomes that are currently publicly available (Table 3).

Utilization of the pentose, xylose, by *L. plantarum* and *L. pentosus* has been discussed in the literature given the use of such a trait to establish *L. pentosus* as a species apart from *L. plantarum* (46). In general LAB utilize pentose sugars, including xylose, via the Pentose Phosphate / Glycolytic Pathway or the Phosphoketolase Pathway (47, 48, 49). In this study, putative genes coding for a Xylose Symporter (*xylR*), D-Xylose-5-Phosphate 3-Epimerase and the Xylulokinase were frequently detected in the *L. buchneri* genome sequences but seldom found in the *L. plantarum* genomes (Table 3). The genomes of several strains of *L. pentosus*, *P. pentosaceus*, and *L. brevis* also harbored some of the putative genes (Table 3). Additionally, genes putatively encoding for key enzymes in the Pentose Phosphate Pathway were found in the *L. plantarum* and *L. pentosus* genome sequences studied which convert D-Ribulose-5-Phosphate to D-Glyceraldehyde-3-Phosphate, an important Glycolysis intermediary. In this study, neither *L. plantarum* ATCC 14917, WCSF1 and 3.2.8 nor *L. pentosus* LA0445, ATCC 8041 and 1.8.9 were energized by xylose in FCJM at pH 4.7 ± 0.1 (Table 4). However, *L. brevis* ATCC14869, ATCC8287 and 7.2.43 converted xylose to lactic acid and acetic acid deriving energy to double (Table 4). It has been reported that strains of *L. brevis* and *L. plantarum* are able to ferment xylose via the Phosphoketolase Pathway (24). Specific strains of *P. pentosaceus* and *L. pentosus* were found to metabolize xylose, but not *L. buchneri* and *L. plantarum* by another group (19, 20, 21, 22, 23). The *L. buchneri* genome sequences were found to be severely impaired with regards to putative genes coding for enzymes involved in xylose utilization and the end of glycolysis (Table 3). However, *L. buchneri* ATCC4005, LA1147 and LA1149 utilized nearly all the xylose supplemented in the FCJM (18.65 ± 0.49) converting it to lactic acid and acetic acid, dropping the pH from 4.7 ± 0.1 to about 4.2 and deriving energy for growth to 8 log CFU/mL (Table 4). Thus, it was apparent that the ability of specific LAB strains to utilize xylose is dependent on their niche and/or culture conditions.

The ability of *L. buchneri* to utilize L-citrulline under conditions similar to those present in commercial cucumber fermentations after glucose and fructose are consumed was confirmed in this study. Genes coding for the L-citrulline-aspartate ligase and the arginine-succinate lyase were commonly found in the *L. plantarum*, *L. pentosus*, and *L. buchneri* genome sequences, but not in the *L. brevis* and *P. pentosaceous* genome sequences (Table 3). All the *P. pentosaceus* and *L. buchneri* genome sequences and more than 85% of the *L. brevis* genomic sequences were unique in encoding for a putative arginine deiminase which interconverts L-citrulline to arginine (Table 3). Arginine deiminase is used by LAB to convert arginine into L-citrulline as an intermediate and then to ammonia, ornithine, ATP, and CO_2_ (35, 50, 30, 51, 50, 30). However, *L. buchneri* was unique in utilizing L-citrulline and deriving energy for growth with minimal changes in pH (Table 4). This observation is consistent with those made by others from wine fermentations (51, 50, 30). An increase in the FCJM L-citrulline concentration was measured from samples inoculated with *L. plantarum*, *L. pentosus* and *L. brevis* with significant decreases in pH and increases in colony counts, suggesting the conversion of arginine naturally present in the medium to the non-proteinaceous amino acid (Table 4). Several studies focusing on wine fermentations showed that a strain of *L. buchneri* (CUC-3) can metabolize either arginine or L-citrulline using the ADI pathway (50, 30). Some strains of *L. buchneri, L. brevis, L. hilgardii,* and *P. pentosaceus* can synthesize L-citrulline from the degradation of arginine in wine fermentations (51). Two strains of *L. plantarum* (N8 and N4) have been found capable of utilizing both L-citrulline and arginine in a stressful environment such asorange juice (52). On the other hand, L-citrulline accumulation occurs in soy sauce fermentation during the lactic acid production stage and the alcoholic fermentation (53). Strains of *Bacillus amyloliquefaciens* are able to metabolize L-citrulline and ethyl carbamate in soy sauce fermentation (53, 24).

An additional experiment was conducted to confirm the ability of *L. buchneri* to utilize L-citrulline in the presence of limiting and excess sugars at pH 4.7 and 3.7. A mixed *L. buchneri* inocula consisting of strains LA0030, LA1149, and LA1147, was used to inoculate FCJM supplemented with L-citrulline, glucose or a combination of the two (Table 5). L-citrulline was utilized in the presence of limiting glucose and converted to ammonia and ornithine inducing an increase in pH (Table 5). The presence of glucose enable the conversion of L-citrulline into an unidentified product, which is presumed to be arginine that had been incorporated into biomass or other metabolic activity, but not to ammonia or ornithine (Table 5) (35, 33, 30). Production of ammonia and ornithine by *L. buchneri* was also observed in the unsupplemented FCJM suggesting that arginine had been utilized as a source of L-citrulline (Table 5). The presence of L-citrulline in the FCJM enhanced growth of *L. buchneri* as compared to the unsupplemented control and proliferation of the LAB at the lower pH (3.7 ± 0.1) (Table 5).

Table 6 demonstrates inoculation of a mixed starter cultureof *L. brevis* and *L. buchneri* enables a more complete fermentation in FrCJ medium supplemented with xylose, trehalose and L-citrulline as compared to the use of *L. pentosus* in the starter culture. *L. brevis* was able to utilize about 50% of the xylose supplemented in FrCJ medium and a substantial portionof the trehalose and L-citrulline. Co-inoculation of *L. brevis* and *L. buchneri* in the FrCJ medium resulted in the removal of all the three alternate energy sources supplemented, the highest production of lactic and acetic acids and an ending pH higher than that observed when *L. pentosus* was inoculated. These observations suggest that the utilization of L-citrulline could have raised the pH enabling more acid production and a higher end of fermentation pH. Additionally, diversion of the sugars to acetic acid instead of lactic acid could have contributed to a higher final pH. Acetic acid has a higher dissociation constant as compared to lactic acid. Interestingly, no changes in pH were observed after the primary fermentation was completed by *L. brevis* and *L. buchneri* suggesting microbial stability for about 50 days under anaerobiosis (Table 6).

Utilization of a mixed starter culture of *L. pentosus*, *L. brevis* and *L. buchneri* in FrCJ medium resulted in the removal of L-citrulline immediately after the conversion of sugars to lactic acid and acetic acid, a slightly higher pH as compared to cultures inoculated with *L. brevis* and *L. pentosus* and stable colony counts from MRS agar plates (Table 7). The use of a mixed culture of *L. pentosus* and *L. brevis* resulted in the presence of L-citrulline in the FrCJ medium after the primary fermentation was concluded (Table 7). Utilization of *L. brevis* alone resulted in the partial utilization of xylose and an incomplete fermentation. The use of a tripartite starter culture for the fermentation of cucumber is a viable strategy to prevent spoilage of fermented cucumbers during bulk storage that merits further investigation.

It is concluded that the occasional presence of trehalose and xylose in commercial cucumber fermentations does not represent a steady alternate energy source for spoilage organisms given that the sugars can be metabolized by the LAB prevailing in the system, *L. pentosus*, *L. plantarum* and *L. brevis*. The presence of L-citrulline in commercial cucumber fermentations at a pH of 3.7 or above could propel the development of spoilage during bulk storage given that it is not utilized by *L. pentosus*, *L. plantarum* and *L. brevis* during primary fermentation. Utilization of a combination of *L. pentosus*, *L. brevis* and *L. buchneri* as a starter culture in FrCJ medium resulted in the early removal of alternate energy sources such as xylose, trehalose and L-citrulline. Further studies are needed to determine if the application of the tripartite starter culture proposed here could complete a cucumber fermentation and generate a microbiologically stable fermented product for bulk storage.

## Acknowledgments

The authors thank Ms. Sandra Parker and Ms. Janet Hayes and Mr. Robert Price with the USDA-ARS Food Science Research Unit located in Raleigh, NC for administrative and technical support, respectively. The authors also thank the Turkish Government for the fellowship support for Redife Aslihan Ucar. No conflict of interest is declared.

## References

1. Handley, L.W. Pharr, D.M., McFeeters, R.F., 1983. Relationship between galactinol synthase activity and sugar composition of leaves and seeds. J. Am. Soc. Hortic. Sci., 108, 600–605.

2. McFeeters, R.F., 1992. Cell wall monosaccharide changes during softening of brined cucumber mesocarp tissue. J. Food Sci., 57(4), 937–940.

3. Pharr, D.M., Sox, H.N., Smart, E.L., Lower, R.L., Fleming, H.P., 1977. Identification and distribution of soluble saccharides in pickling cucumber plants and their fate in fermentation. J. Am. Soc. Hortic. Sci. 102(4), 406–409.

4. Johanningsmeier, S.D., McFeeters, R.F., 2015. Metabolic footprinting of *Lactobacillus buchneri* strain LA1147 during anaerobic spoilage of fermented cucumbers. Int. J. Food Microbiol., 215, 40–48.

5. Yu, J., Gao, W., Qing, M., Sun, Z., Wang, W., Liu, W., Pan, L., Sun, T., Wang, H., Bai, N., Zhang, H., 2012. Identification and characterization of lactic acid bacteria isolated from traditional pickles in Sichuan, China. J. Gen. Appl. Microbiol., 58(3), 163–172.

6. Tamang, J.P., Tamang, B., Schillinger, U., Franz, C.M., Gores, M., Holzapfel, W.H., 2005. Identification of predominant lactic acid bacteria isolated from traditionally fermented vegetable products of the Eastern Himalayas. Int. J. Food Microbiol., 105(3), 347–356.

7. Kleerebezem, M., Hugenholtz, J., 2003. Metabolic pathway engineering in lactic acid bacteria. Curr. Opin. Biotechnol., 14(2), 232–237.

8. Lorca, G.L., Barabote, R.D., Zlotopolski, V., Tran, C., Winnen, B., Hvorup, R.N., Stonestrom, A.J., Nguyen, E., Huang, L.W., Kim, D.S., Saier Jr, M.H., 2007. Transport capabilities of eleven gram-positive bacteria: comparative genomic analyses. Biochim. Biophys. Acta, Biomembr., 1768(6), 1342–1366.

9. Siezen, R.J., van Hylckama Vlieg, J.E., 2011. Genomic diversity and versatility of *Lactobacillus plantarum*, a natural metabolic engineer. In Microbial cell factories. BioMed. Cent., 10(1), S3.

10. Westby, A., Nuraida, L., Owens, J.D., Gibbs, P.A., 1993. Inability of *Lactobacillus plantarum* and other lactic acid bacteria to grow on D-ribose as sole source of fermentable carbohydrate. J. Appl. Bacteriol., 75(2), 168–175.

11. Siezen, R.J., Tzeneva, V.A., Castioni, A., Wels, M., Phan, H.T., Rademaker, J.L., Starrenburg, M.J., Kleerebezem, M., Molenaar, D., Van Hylckama Vlieg, J.E., 2010. Phenotypic and genomic diversity of *Lactobacillus plantarum* strains isolated from various environmental niches. Environ. Microbiol., 12(3), 758–773.

12. Sanders, J.W., Oomes, S.J.C.M., Membré, J.M., Wegkamp, A., Wels, M., 2015. Biodiversity of spoilage lactobacilli: phenotypic characterisation. Food Microbiol., 45, 34–44.

13. Ucar, R. A., Pérez-Díaz, I. M., Dean, L. L. Submitted. Gentiobiose and cellobiose content in fresh and fermenting cucumbers and utilization of such disaccharides by lactic acid bacteria in fermented cucumber juice medium. J. Food Sci.

14. Crowe, J.H., Crowe, L.M., Oliver, A.E., Tsvetkova, N., Wolkers, W., Tablin, F., 2001. The trehalose myth revisited: introduction to a symposium on stabilization of cells in the dry state. Cryobiology., 43(2), 89–105.

15. Romero, C., Bellés, J.M., Vayá, J.L., Serrano, R., Culiáñez-Macià, F.A., 1997. Expression of the yeast trehalose-6-phosphate synthase gene in transgenic tobacco plants: Pleiotropic phenotypes include drought tolerance. Planta., 201(3), 293–297.

16. Goddijn, O.J., van Dun, K., 1999. Trehalose metabolism in plants. Trends Plant Sci., 4(8), 315–319.

17. Leslie, S.B., Israeli, E., Lighthart, B., Crowe, J.H., Crowe, L.M., 1995. Trehalose and sucrose protect both membranes and proteins in intact bacteria during drying. Appl. Environ. Microbiol., 61(10), 3592–3597.

18. Colaço, C.A.L.S., Smith, C.J.S., Sen, S., Roser, D.H., Newman, Y., Ring, S., Roser, B.J., 1994. Chemistry of protein stabilization by trehalose. In ACS symposium series, 222–240. DOI: 10.1021/bk-1994-0567.ch014.

19. Bringel, F., Curk, M.C., Hubert, J.C., 1996. Characterization of lactobacilli by Southern-type hybridization with a *Lactobacillus plantarum* pyrDFE probe. Int. J. Syst. Evol. Microbiol., 46(2), 588–594.

20. Bustos, G., Moldes, A.B., Cruz, J.M., Domínguez, J.M., 2005. Influence of the metabolism pathway on lactic acid production from hemicellulosic trimming vine shoots hydrolyzates using *Lactobacillus pentosus*. Biotechnol. Prog., 21(3), 793–798.

21. Carr, F.J., Chill, D., Maida, N., 2002. The lactic acid bacteria: A literature survey. Crit. Rev. Microbiol. 28(4), 281–370.

22. Chaillou, S., Pouwels, P.H., Postma, P.W., 1999. Transport of d-Xylose in *Lactobacillus pentosus*, *Lactobacillus casei*, and *Lactobacillus plantarum*: Evidence for a mechanism of facilitated diffusion via the Phosphoenolpyruvate: Mannose Phosphotransferase System. J. Bacteriol. 181(16), 4768–4773.

23. Lokman, B.C., Leer, R.J., van Sorge, R., Pouwels, P.H., 1994. Promotor analysis and transcriptional regulation of *Lactobacillus pentosus* genes involved in xylose catabolism. Mol. Gen. Genet., 245(1), 117–125.

24. Zhang, Y., Zeng, F., Hohn, K., Vadlani, P.V., 2016. Metabolic flux analysis of carbon balance in *Lactobacillus* strains. Biotechnol. Prog., 32(6), 1397–1403.

25. Kim, J.H., Shoemaker, S.P., Mills, D.A., 2009. Relaxed control of sugar utilization in *Lactobacillus brevis*. Microbiol., 155(4), 1351–1359.

26. Hammes, W.P., Hertel, C., 2015. Lactobacillus. Bergey’s Manual of Systematics of Archaea and Bacteria, 1–76.

27. Fragkos, K., Forbess, A., 2011. Was citrulline first a laxative substance? The truth about modern citrulline and its isolation. Nihon ishigaku zasshi/J. Jpn. Hist. Med., 57(3), 275–292.

28. Fish, W.W., 2012. A reliable methodology for quantitative extraction of fruit and vegetable physiological amino acids and their subsequent analysis with commonly available HPLC systems. Food Nutr. Sci., 3(06), 863.

29. Yokota, A., Kawasaki, S., Iwano, M., Nakamura, C., Miyake, C., Akashi, K., 2002. Citrulline and DRIP-1 protein (ArgE homologue) in drought tolerance of wild watermelon. Ann. Bot., 89(7), 825–832.

30. Liu, S.Q., Pritchard, G.G., Hardman, M.J., Pilone, G.J., 1996. Arginine catabolism in wine lactic acid bacteria: Is it via the arginine deiminase pathway or the arginase-urease pathway? J. Appl. Bacteriol., 81(5), 486–492.

31. Spano, G., Beneduce, L., Tarantino, D., Giammanco, G.M., Massa, S., 2002. Preliminary characterization of wine lactobacilli able to degrade arginine. World J. Microbiol. Biotechnol., 18(9), 821–825.

32. Arena, M.E., Manca de Nadra, M.C., 2002. Comparative survey in *Lactobacillus plantarum* of the growth and metabolism of arginine and citrulline in different media. J. Agric. Food Chem., 50(22), 6497–6500.

33. Gänzle, M.G., 2015. Lactic metabolism revisited: metabolism of lactic acid bacteria in food fermentations and food spoilage. Curr. Opin. Food Sci., 2, 106–117.

34. Gänzle, M.G., Vermeulen, N., Vogel, R.F., 2007. Carbohydrate, peptide and lipid metabolism of lactic acid bacteria in sourdough. Food Microbiol., 24(2), 128–138.

35. Bauer, R., Dicks, L.M., 2017. Control of malolactic fermentation in wine. A review. S. Afr. J. Enol. Vitic., 25(2), 74–88.

36. Pattee, H.E., Isleib, T.G., Giesbrecht, F.G., McFeeters, R.F., 2000. Investigations into genotypic variations of peanut carbohydrates. J. Agric. Food Chem., 48(3), 750–756.

37. Otaka, S., 2013. Hitachi High Technologies, Dallas, TX. Personal Communication.

38. Chen, I.M., Markowitz, V.M., Chu, K., Palaniappan, K., Szeto, E., Pillay, M., Ratner, A., Huang, J., Andersen, E., Huntemann, M. and Varghese, N., 2016. IMG/M: integrated genome and metagenome comparative data analysis system. Nucl. Acids Res. 45:D1, D507–D516.

39. McFeeters, R.F., Barish, A.O., 2003. Sulfite analysis of fruits and vegetables by high-performance liquid chromatography (HPLC) with ultraviolet spectrophotometric detection. J. Agric. Food Chem., 51(6), 1513–1517.

40. Pérez-Díaz, I.M., Breidt, F., Buescher, R.W., Arroyo-López, F.N., Jiménez-Dıaz, R., Fernandez, A.G., Gallego, J.B., Yoon, S.S., Johanningsmeier, S.D., 2016. Fermented and acidified vegetables. In Compendium of methods for the microbiological examination of foods, 4th edn. pp. 521–532. Washington, D.C.: American Public Health Association.

41. Gänzle, M., Follador, R., 2012. Metabolism of oligosaccharides and starch in *lactobacilli*: A review. Front. Microbiol., 3, 340.

42. Andersson, U., Molenaar, D., Rådström, P., de Vos, W.M., 2005. Unity in organisation and regulation of catabolic operons in *Lactobacillus plantarum*, *Lactococcus lactis* and *Listeria monocytogenes*. Syst. Appl. Microbiol., 28(3), 187–195.

43. Francl, A.L., Thongaram, T., Miller, M.J., 2010. The PTS transporters of *Lactobacillus gasseri* ATCC 33323. BMC Microbiol., 10(1), 77.

44. Mao, Y., Chen, M., Horvath, P., 2015. *Lactobacillus herbarum* sp. nov., a species related to *Lactobacillus plantarum*. Int. J. Syst. Evol. Microbiol., 65(12), 4682–4688.

45. Sterr, Y., Weiss, A., Schmidt, H., 2009. Evaluation of lactic acid bacteria for sourdough fermentation of amaranth. Int. J. Food Microbiol., 136(1), 75–82.

46. Fred, E.B., Peterson, W.H., Anderson, J.A., 1921. The characteristics of certain pentose destroying bacteria, especially as concerns their action on arabinose and xylose. J. Bio. Chem., 48(2), 385–412.

47. Abdel-Rahman, M.A., Tashiro, Y., Zendo, T., Shibata, K., Sonomoto, K., 2011. Isolation and characterization of lactic acid bacterium for effective fermentation of cellobiose into optically pure homo L-(+)-lactic acid. Appl. Microbiol. Biotechnol., 89(4), 1039–1049.

48. Okano, K., Yoshida, S., Yamada, R., Tanaka, T., Ogino, C., Fukuda, H., Kondo, A., 2009. Improved production of homo-D-lactic acid via xylose fermentation by introduction of xylose assimilation genes and redirection of the phosphoketolase pathway to the pentose phosphate pathway in L-lactate dehydrogenase gene-deficient *Lactobacillus plantarum*. Appl. Environ. Microbiol., 75(24), 7858–7861.

49. Tanaka, K., Komiyama, A., Sonomoto, K., Ishizaki, A., Hall, S., Stanbury, P., 2002. Two different pathways for D-xylose metabolism and the effect of xylose concentration on the yield coefficient of L-lactate in mixed-acid fermentation by the lactic acid bacterium *Lactococcus lactis* IO-1. Appl. Microbiol. Biotechnol., 60(1-2), 160–167.

50. Liu, S.Q., Davis, C.R., Brooks, J.D., 1995. Growth and metabolism of selected lactic acid bacteria in synthetic wine. Am. J. Enol. Vitic., 46(2), 166–174.

51. Araque, I., Reguant, C., Rozès, N., Bordons, A., 2011. Influence of wine-like conditions on arginine utilization by lactic acid bacteria. Int. Microbiol., 14, 225–233.

52. Arena, M.E., Saguir, F.M., de Nadra, M.M., 1999. Arginine dihydrolase pathway in *Lactobacillus plantarum* from orange. Int. J. Food Microbiol., 47(3), 203–209.

53. Fang, F., Zhang, J., Zhou, J., Zhou, Z., Li, T., Lu, L., Zeng, W., Du, G., Chen, J., 2018. Accumulation of citrulline by microbial arginine metabolism during alcoholic fermentation of soy sauce. J. Agric. Food Chem., 66(9), 2108–2113.

54. Hols, P., Slos, P., Dutot, P., Reymund, J., Chabot, P., Delplace, B., Delcour, J., Mercenier, A., 1997. Efficient secretion of the model antigen M6-gp41E in *Lactobacillus plantarum* NCIMB 8826. Microbiol., 143(8), 2733–2741.

55. Pérez-Díaz, I.M., Hayes, J.S., Medina-Pradas, E., Anekella, K., Daughtry, K.V., Dieck, S., Levi, M., Price, R., Butz, N., Lu, Z., Azcarate-Peril, M., 2017. Reassessment of the succession of lactic acid bacteria in commercial cucumber fermentations and physiological and genomic features associated with their dominance. Food Microbiol., 63, 217–227. DOI: 10.1016/j.fm.2016.11.025.

56. Fleming, H.P., McFeeters, R.F., Daeschel, M.A., Humphries, E.G., Thompson, R.L., 1988. Fermentation of cucumbers in anaerobic tanks. J. Food Sci., 53(1), 127–133.

57. Dunn, M.S., Shankman, S., Camien, M.N., Block, H., 1947. The amino acid requirements of twenty-three lactic acid bacteria. J. Bio. Chem., 168(1), 1–22.

58. Rogosa, M., Hansen, P.A., 1971. Nomenclatural considerations of certain species of *Lactobacillus* Beijerinck. Int. J. Syst. Evol. Microbiol., 21(2), 177–186.

59. Franco, W., Pérez-Díaz, I.M., Johanningsmeier, S.D., McFeeters, R.F., 2012. Characteristics of spoilage-associated secondary cucumber fermentation. Appl. Environ. Microbiol., 78(4), 1273–1284.

